# A lever hypothesis for Synaptotagmin-1 action in neurotransmitter release

**DOI:** 10.1101/2024.06.17.599417

**Authors:** Klaudia Jaczynska, Victoria Esser, Junjie Xu, Levent Sari, Milo M. Lin, Christian Rosenmund, Josep Rizo

## Abstract

Neurotransmitter release is triggered in microseconds by Ca^2+^-binding to the Synaptotagmin-1 C_2_ domains and by SNARE complexes that form four-helix bundles between synaptic vesicles and plasma membranes, but the coupling mechanism between Ca^2+^-sensing and membrane fusion is unknown. Release requires extension of SNARE helices into juxtamembrane linkers that precede transmembrane regions (linker zippering) and binding of the Synaptotagmin-1 C_2_B domain to SNARE complexes through a ‘primary interface’ comprising two regions (I and II). The Synaptotagmin-1 Ca^2+^-binding loops were believed to accelerate membrane fusion by inducing membrane curvature, perturbing lipid bilayers or helping bridge the membranes, but SNARE complex binding orients the Ca^2+^-binding loops away from the fusion site, hindering these putative activities. Molecular dynamics simulations now suggest that Synaptotagmin-1 C_2_ domains near the site of fusion hinder SNARE action, providing an explanation for this paradox and arguing against previous models of Sytnaptotagmin-1 action. NMR experiments reveal that binding of C_2_B domain arginines to SNARE acidic residues at region II remains after disruption of region I. These results and fluorescence resonance energy transfer assays, together with previous data, suggest that Ca^2+^ causes reorientation of the C_2_B domain on the membrane and dissociation from the SNAREs at region I but not region II. Based on these results and molecular modeling, we propose that Synaptotagmin-1 acts as a lever that pulls the SNARE complex when Ca^2+^ causes reorientation of the C_2_B domain, facilitating linker zippering and fast membrane fusion. This hypothesis is supported by the electrophysiological data described in the accompanying paper.

**Significance statement:** Neurotransmitter release requires SNARE complexes that fuse synaptic vesicles with the plasma membrane and the Ca^2+^-sensor synaptotagmin-1, which was thought to facilitate membrane fusion directly through its Ca^2+^-binding loops. However, binding of Synaptotagmin-1 to SNARE complexes orients these loops away from the fusion site. Using molecular dynamics simulations, we show that placing Synaptotagmin-1 at the fusion site hinders the action of SNARE complexes. Spectroscopic studies show that Ca^2+^ binding to Synaptotagmin-1 can change its interactions with SNARE complexes and, together with molecular modeling, suggest that Synaptotagmin-1 acts as a lever, pulling SNARE complexes and thus facilitating their action on the membranes to induce fusion. Functional studies described in the accompanying paper support this hypothesis.

---

Release of neurotransmitters by Ca^2+^-triggered synaptic vesicle exocytosis is crucial for neuronal communication. Exocytosis involves several steps, including vesicle tethering to presynaptic active zones, priming of the vesicles to a release-ready state(s) and vesicle fusion with the plasma membrane, which occurs very fast upon Ca^2+^ influx into a presynaptic terminal (1) [less than 60 μs in fast synapses (2)]. The basic steps that lead to exocytosis have been reconstituted with the main components of the neurotransmitter release machinery (3-6) and the functions of these components have been defined (7-9). The SNAP receptors (SNAREs) syntaxin-1, SNAP-25 and synaptobrevin form tight complexes (10) that consist of four-helix bundles (11, 12) and bring the membranes together as they assemble (zipper) from the N- to the C-terminus (13), likely inducing membrane fusion (14) by promoting lipid acyl chain encounters at the polar membrane-membrane interface (15). N-ethylmaleimide sensitive factor (NSF) and soluble NSF attachment proteins (SNAPs) disassemble SNARE complexes (10) to recycle the SNAREs (16). Munc18-1 and Munc13s organize SNARE complex formation by an NSF-SNAP-resistant mechanism (3, 17) in which Munc18-1 binds first to a closed conformation of syntaxin-1 (18, 19) and later binds to synaptobrevin, forming a template complex (20-22) for SNARE assembly while Munc13 bridges the two membranes (4, 23) and opens syntaxin-1 (24). In the resulting primed state, the SNARE complex is bound to Synaptotagmin-1 (Syt1) (25) and to complexin (26), forming a spring-loaded macromolecular assembly (27) that prevents premature fusion but is ready to trigger fast fusion when Ca^2+^ binds to Syt1 (28).

The cytoplasmic region of Syt1 is formed mostly by two C_2_ domains (C_2_A and C_2_B) that bind three and two Ca^2+^ ions, respectively, via loops at the tip of β-sandwich structures (29-31). While it is well established that Syt1 triggers neurotransmitter release through its C_2_ domains (28, 32), and that Ca^2+^- binding to the C_2_B domain is particularly crucial for release (33, 34), the underlying mechanism remains highly enigmatic. Ca^2+^ induces insertion of these loops into membranes (35, 36), which was proposed to facilitate membrane fusion by perturbing the bilayers (35, 37, 38) and/or inducing membrane curvature (39, 40). In addition, the Syt1 C_2_B domain binds to phosphatidylinositol 4,5-bisphosphate (PIP_2_) through a polybasic region on the side of the β-sandwich (41) and was proposed to help induce fusion by bridging the two membranes (42, 43). Many studies reported diverse interactions of Syt1 with the SNAREs that were generally enhanced by Ca^2+^ [reviewed in (44, 45)]. However, a fragment spanning the two Syt1 C_2_ domains (C_2_AB) binds with higher affinity to nanodiscs containing SNARE complex than to plain nanodiscs in the absence of Ca^2+^ while the affinity is similar in the presence of Ca^2+^, suggesting that Ca^2+^ actually induces dissociation of C_2_AB from membrane-anchored SNARE complexes (46, 47) or at least weakens the C_2_AB-SNARE interactions.

Three structures of Syt1-SNARE complexes have been described, each revealing binding through distinct surfaces of the C_2_B domain that are not involved in Ca^2+^ binding (25, 48, 49). However, strong evidence suggests that, among these three, the key functionally relevant binding mode is mediated by a so-called primary interface (25, 47, 50) formed by two regions of C_2_B: region I involving E295 and Y338; and region II involving R281, R398 and R399. Evidence suggested that this interface is important for vesicle priming (49, 51), which is strongly supported by the accompanying paper (52) and likely arises because Syt1 cooperates with Munc18-1 and Munc13-1 in mediating Ca^2+^-independent assembly of trans-SNARE complexes (17). Intriguingly, while an R398Q,R399Q mutation in region II that disrupts C_2_B-SNARE complex binding (47) strongly impairs neurotransmitter release (25, 53), release is also impaired by a E295A,Y338W mutation in region I (25) that enhances binding (47). Moreover, this binding mode orients the C_2_B Ca^2+^- binding loops away from the site of membrane fusion, which hinders a direct role for the loops in fusion. Note also that Ca^2+^-dependent binding to PIP_2_-containing membranes induces an approximately perpendicular orientation of the C_2_B domain with respect to the bilayer (54) that is incompatible with SNARE complex binding via the primary interface (47). These findings suggested that binding of Syt1 to the SNARE complex via the primary interface is important to generate the primed state but hinders membrane fusion, and Ca^2+^-induced dissociation of Syt1 from the SNAREs relieves the inhibition (47).

While this model explains a large amount of data and release-of-inhibition models have been popular since 30 years ago (55), there is evidence that Syt1 is not merely a SNARE inhibitor and exerts an active action to trigger release upon Ca^2+^ binding (7, 44). Dissociation from the SNAREs might allow the Syt1 C_2_ domains to reorient toward the site of fusion such that they can directly facilitate fusion through its Ca^2+^-binding loops (47), but this notion does not bode well for the fast speed of release and Ca^2+^- induced dissociation of Syt1 from membrane-anchored SNARE complexed has not been demonstrated. Moreover, it is not clear how actions of Syt1 that have been proposed to facilitate membrane fusion such as bilayer perturbation (35, 37, 38), induction of membrane curvature (39, 40) or membrane bridging (42, 43) can cooperate with the SNAREs to induce fast membrane fusion, particularly after molecular dynamics (MD) simulations revealed a natural mechanism for SNARE complexes to induce fast, microsecond-scale membrane fusion (15). In this mechanism, extension of the synaptobrevin and syntaxin-1 helices that form the four-helix bundle into the juxtamembrane (jxt) linkers that precede the transmembrane (TM) regions (referred to as jxt linker zippering) is key to catalyze encounters between the lipid acyl chains of the two bilayers that initiate fusion, but it is unclear how the Syt1 Ca^2+^-binding loops can facilitate these events.

To address these questions and shed light into how Syt1 triggers neurotransmitter release, we have used a combination of all-atom MD simulations with NMR and fluorescence spectroscopy assays. The simulations suggest that placing the Syt1 C_2_ domains near the site of fusion severely hinders the ability of SNARE complexes to bring the membranes together and induce membrane fusion, which provides an explanation for why the primary interface orients the C_2_B Ca^2+^-binding loops away from the site of fusion and argues against the notion that Syt1 facilitates membrane fusion by inducing membrane curvature, perturbing the bilayers or bridging the membranes. NMR experiments show that binding of the Syt1 C_2_B domain to the SNARE complex via the arginines of region II of the primary interface remains when binding through region I is disrupted. Fluorescence resonance energy transfer (FRET) show that Ca^2+^ does not dissociate a fragment spanning the two C_2_ domains of Syt1 (C_2_AB) from membrane-anchored SNARE complexes but induces a reorientation of the C_2_B domain with respect to the SNARE four-helix bundle. These results, together with previous electron paramagnetic resonance (EPR) data (54), suggest a model whereby Ca^2+^ binding causes dissociation of region I but not region II of the primary interface and reorientation of C_2_B on the membrane, which pulls the SNARE complex through ionic interactions and facilitates linker zippering to initiate fast membrane fusion. While further research will be needed to test this lever hypothesis of Syt1 action, the central notion that neurotransmitter release is triggered by Ca^2+^- induced re-modeling of the Syt1-SNARE primary interface is strongly supported by multiple correlations between our NMR data and electrophysiological results described in the accompanying paper (52).

## Results

### Syt1 C_2_ domains close to the fusion site hinder SNARE action

Since all-atom MD simulations have provided a powerful tool to visualize the neurotransmitter release machinery bridging two membranes (27, 56) and understand how the SNAREs mediate membrane fusion (15), we carried out several all-atom MD simulations to investigate the merits of various models that have been proposed for how Syt1 accelerates membrane fusion. Similar to our previous studies (15, 27), the systems used for all simulations contained four trans-SNARE complexes bridging a vesicle and a flat bilayer with lipid compositions that resemble those of synaptic vesicles and synaptic plasma membranes, respectively (57, 58), with or without Syt1 C_2_AB molecules (Table S1). Complexin was not included in these simulations for simplicity, as fast neurotransmitter release is impaired but not abolished in the absence of complexins (59) and we wanted to focus on how the functions of Syt1 and the SNAREs are coupled.

In a first simulation, we used a system of four trans-SNARE complexes that were zippered to distinct extents at the C-terminus, and included four C_2_AB molecules bound to five Ca^2+^ ions each and with the Ca^2+^-binding loops pointing toward the flat bilayer without contacting it (Fig. S1A) to investigate whether the Syt1 C_2_ domains spontaneously insert into the bilayers, perturb or bridge the bilayers, and perhaps even initiate membrane fusion (referred to as cac2absc simulation). The SNARE complexes were in the same configurations and positions as those of a simulation of the primed state described previously [prsg simulation in (27)], with different extents of assembly. During the 668 ns of this simulation, the two membranes were brought into contact (Fig. 1A), but there was no substantial progress in zippering of the SNARE complexes and the C_2_ domains of the distinct C_2_AB molecules exhibited different orientations with respect to each other, to the SNARE complexes and to the membranes (Fig. S1B,C). One of the C_2_B domains adopted an approximately perpendicular orientation with respect to the flat bilayer, inserting the Ca^2+^- binding loops into this bilayer and binding to the vesicle through the opposite end of the domain that contains R398 and R399 (Fig. 2A), as predicted in membrane-bridging models of Syt1 function (42, 43). However, the Ca^2+^-binding loops inserted only to a small extent into the acyl region of the bilayer, and there was no overt perturbation of the bilayer or induction of curvature (Fig. 2A). The other C_2_ domains adopted more slanted or even parallel orientations with respect to the flat bilayer (Fig. S1B,C). One of the C_2_B domains had the Ca^2+^-binding loops oriented toward the center of the membrane-membrane interface and was located next to the C-terminus of a SNARE complex that was almost fully assembled from the beginning of the simulation (Fig. S1D), which could facilitate cooperation of the SNAREs and the C_2_B domain in inducing membrane fusion. We note however that this configuration was already present early in the simulation (at 180 ns), but there was no initiation of fusion during the rest of the simulation.

**Figure 1.**
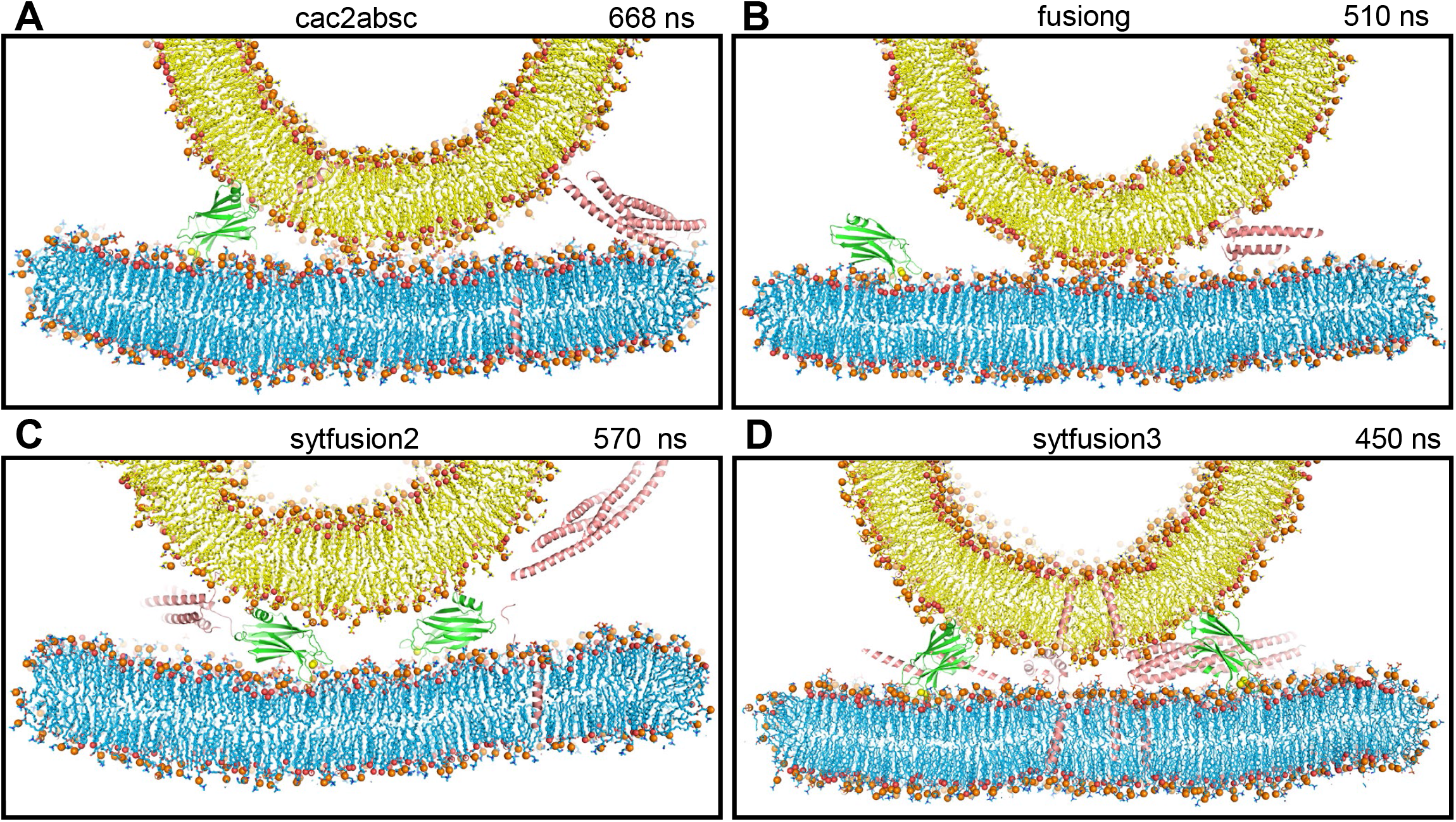
The Syt1 C_2_ domains hinder SNARE action when located near the site of fusion. (*A-D*) Diagrams showing thin slices of frames taken at 668 ns of the cac2absc simulation (*A*), 510 ns of the fusiong simulation (*B*), 570 ns of the sytfusion2g simulation (*C*) and 450 ns of the sytfusion3 simulation (*D*). Lipids are shown as stick models with nitrogen atoms in dark blue, oxygens in red, phosphorus in orange and carbon atoms in yellow (vesicle) or light blue (flat bilayer). Phosphorous atoms of phospholipids and the oxygen atoms of cholesterol molecules are shown as spheres to illustrate the approximate locations of lipid head groups. Proteins are represented by ribbon diagrams, with SNARE complexes in salmon color, Syt1 C_2_A domain in slate blue and Syt1 C_2_B domain in green. Ca^2+^ ions are shown as yellow spheres. Note that, because the slices shown are thin, only portions of some of the proteins, if any, are seen in the slices. The same color-coding was used in all the figures except when noted otherwise.

**Figure 2.**
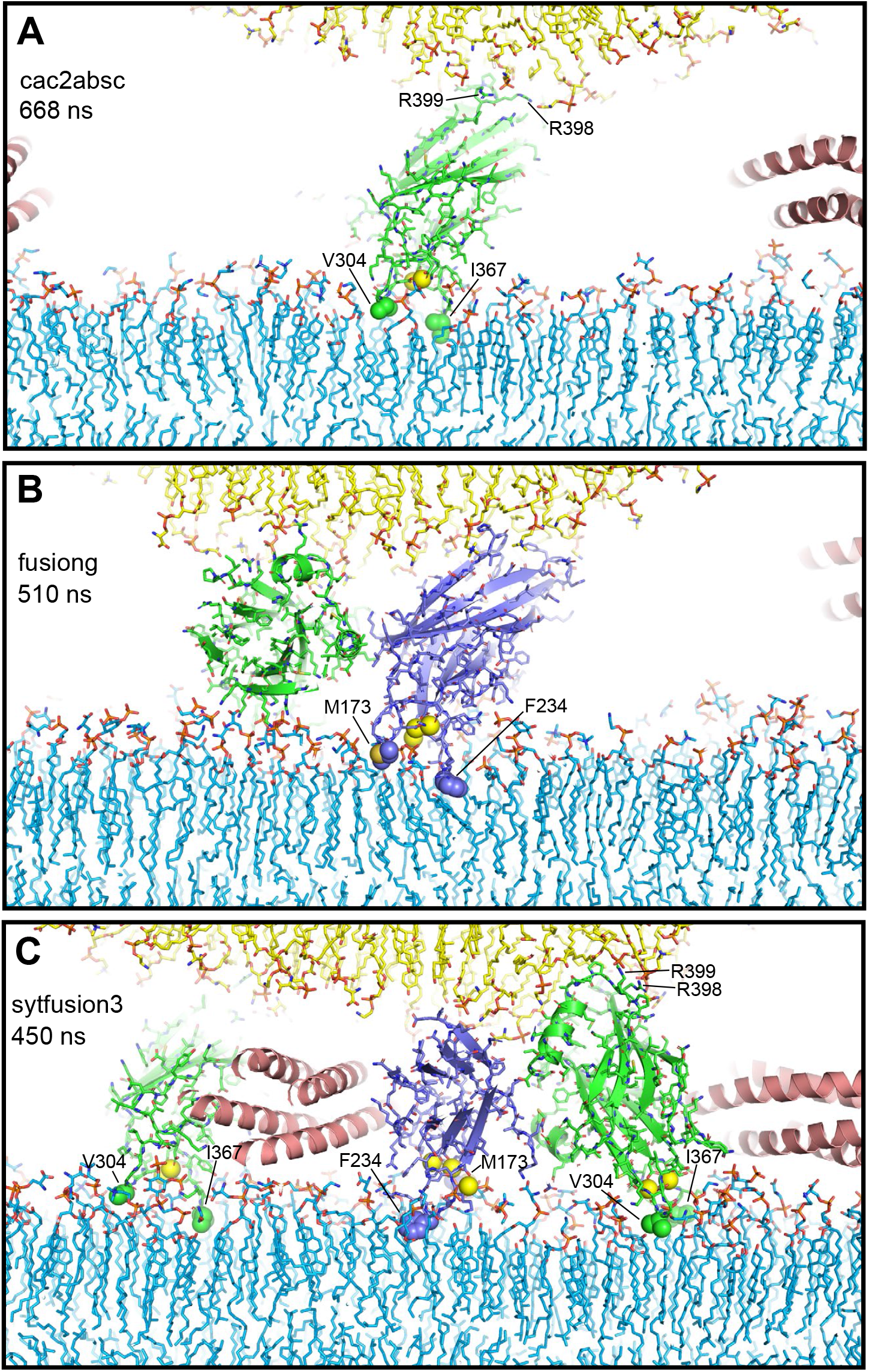
Syt1 C_2_ domain Ca^2+^-binding loops do not insert deeply into lipid bilayers and cause limited bilayer perturbation. The diagrams show examples of C_2_ domains with their Ca^2+^-binding loops interacting with the flat bilayer from frames taken at 668 ns of the cac2absc simulation (*A*), 510 ns of the fusiong simulation (*B*) and 450 ns of the sytfusion3 simulation (*C*). Lipids are shown as stick models. Ca^2+^ ions are shown as yellow spheres. SNARE complexes are represented by ribbon diagrams and Syt1 C_2_ domains by ribbon diagrams and stick models with nitrogen atoms in dark blue, oxygen in red, sulfur in yellow orange and carbon colored in slate blue (C_2_A) and green (C_2_B). Other color coding is as in Fig. 1. The hydrophobic residues at the tips of the Ca^2+^-binding loops that insert into the flat bilayer are shown as spheres and labeled. R398 and R399 at the opposite end of the C_2_B domain are labeled in (*A, C*) to illustrate how the C_2_B domain can bridge two membranes as predicted (42).

We reasoned that the probability of observing membrane fusion in this simulation might have been hindered because three SNARE complexes were partially assembled. Thus, we generated a similar system but with the four SNARE four-helix bundles almost fully assembled and four C_2_AB molecules with the C_2_B domain Ca^2+^-binding loops oriented toward the center of the membrane-membrane interface (fusiong simulation) (Fig. S2A). We performed an MD simulation of this system for 510 ns, but we again did not observe initiation of fusion (Fig. 1B and S2B).

To examine the possibility that the C_2_AB molecules might actually hinder SNARE action in the fusiong simulation, we performed a parallel 510 ns simulation with an identical system that lacked the four C_2_AB molecules (nosytfusion simulation) (Fig. S3A). There was again no fusion in this simulation (Fig. S3B), but the two membranes came into contact more quickly than in the presence of C_2_AB molecules (Fig. S2C and S3C). It is also noteworthy that at the end of the nosytfusion simulation the contact between the two bilayers (Fig. S3D) was considerably more extensive than at the end of the fusiong simulation (Fig. 1B). Such extended interfaces were observed in a previous simulation of a similar system (27) and in cryo-electron microscopy (cryo-EM) images of SNARE-mediated liposome fusion reactions (60). Hence, our results suggest that the C_2_AB molecules of the fusiong simulation hindered the action of the SNAREs in bringing the membranes together and the formation of extended interfaces.

We also explored the notion that the Ca^2+^-binding loops of both Syt1 C_2_ domains might play a direct role in membrane fusion if the C_2_ domains are located between the two membranes with the Ca^2+^- binding loops oriented toward the center of the membrane-membrane interface such that one of the loops can bind to the vesicle and the other to the flat bilayer. In this configuration, Ca^2+^-binding might favor movement of lipids toward the Ca^2+^-binding sites to destabilize the bilayers and initiate fusion. To test this idea, we performed a 570 ns MD simulation of a system analogous to the nosytfusion system but including two C_2_AB molecules between the membranes (sytfusion2g simulation) (Fig. S4A). We did not observe the postulated lipid movements during the simulation and it became clear that the C_2_AB molecules hindered the action of the SNARE complexes in bringing the membranes together (Fig. 1C and S4B).

In these simulations, the SNARE complexes may have been too far from the center of the bilayer-bilayer interface to effectively induce membrane fusion. Placing them closer to the center might facilitate fusion and would prevent the C_2_ domains from coming close to the site of fusion because of steric hindrance. However, the C_2_ domains could be located further from the center, where they could bridge the two membranes as predicted in some models of Syt1 function (42, 43). In such positions, the C_2_ domains might act as wedges that prevent the membranes from coming closer while the SNARE complexes pull the membranes together in the center, resulting in a torque that could help to bend the membranes to initiate fusion [as proposed previously for Munc18-1 function (61)]. To test this model, we used a system with four trans-SNARE complexes closer to the center that we generated previously [fusion2g in (15)], increased the separation between the flat membrane and the vesicle by 1.6 nm to make room for the bridging C_2_ domains, used a restrained simulation to move the syntaxin-1 TM regions to their positions in the translated flat bilayer, and included four C_2_AB molecules in positions in which the two C_2_ domains were poised to bridge the two membranes (Fig. S5A). We ran a 450 ns MD simulation (sytfusion3) and observed that the two membranes came closer to each other but were not brought into contact because of steric hindrance caused by the bridging C_2_ domains, and the SNARE complexes were unable to pull the bilayers together in the center (Fig. 1D and S5B).

The results of all these simulations need to be interpreted with caution because of the limited time of the simulations and hence do not rule out the various models tested. However, the overall results do suggest that C_2_ domains near the site of fusion hinder the action of SNARE complexes in bringing the membranes together. Importantly, throughout these simulations we observed that the hydrophobic residues at the tips of the C_2_ domain Ca^2+^-binding loops often contacted the hydrophobic acyl region of the flat bilayer but did not insert deeply (e.g. Fig. 2A-C), consistent with results from previous MD simulations [e.g. (56)]. Such insertion causes local perturbations but not major alterations of the bilayer structure, and does not induce membrane curvature. Fluorescent probes attached to cysteines in these positions are expected to insert more deeply into the acyl region because they are larger, which has led to the assumption than that the Ca^2+^-binding loops penetrate into the membrane [reviewed in (44, 61)] more than they actually do. Note also that electron microscopy experiments supporting the notion that Syt1 induces membrane curvature used negative stain, which strongly perturbs membranes, and were performed with artificially high protein-to-lipid ratios (1:40) (39, 40) such that proteins should cover most of the lipid surface. All these observations suggest that popular models postulating that Syt1 facilitates membrane fusion by bridging the membranes, perturbing the bilayers or inducing membrane curvature are likely incorrect. Thus, while it seemed paradoxical that binding of the Syt1 C_2_B domain to the SNARE complex via the primary interface orients its Ca^2+^-binding loops away from the fusion site (25), it now appears that this feature makes a lot of sense because it keeps Syt1 away from the fusion site where it would hinder the actions of the SNAREs that induce membrane fusion.

### Ca^2+^ does not cause dissociation of Syt1 C_2_AB from membrane-anchored SNARE complex

The observation that Syt1 C_2_AB binds to nanodiscs containing SNARE complex more tightly to SNARE-free nanodiscs in the absence of Ca^2+^ whereas the affinity is similar in the presence of Ca^2+^ suggested that Ca^2+^ induces dissociation of C_2_AB from membrane-anchored SNARE complex (47), but this notion was not tested directly. To examine whether Ca^2+^ indeed induces such dissociation, we used an approach involving labeling of SNARE complex with bimane on residue 59 of SNAP-25 and replacement of T285 of Syt1 with a tryptophan. This design led to tryptophan-induced quenching of the bimane fluorescence upon binding of Syt1 C_2_B domain to liposome-anchored SC-59-bimane in the absence of Ca^2+^ due to the predicted proximity of the tryptophan to the bimane in the primary interface (62). Correspondingly, we observed strong quenching of the fluorescence of nanodisc-anchored SC-59-bimane upon addition of C_2_AB-T285W in the absence of Ca^2+^ (Fig. 3). Importantly, the quenching remained upon addition of Ca^2+^ (Fig. 3), showing that Ca^2+^ does not induce dissociation of Syt1 C_2_AB from membrane-anchored SNARE complex under the conditions of these experiments.

**Figure 3.**
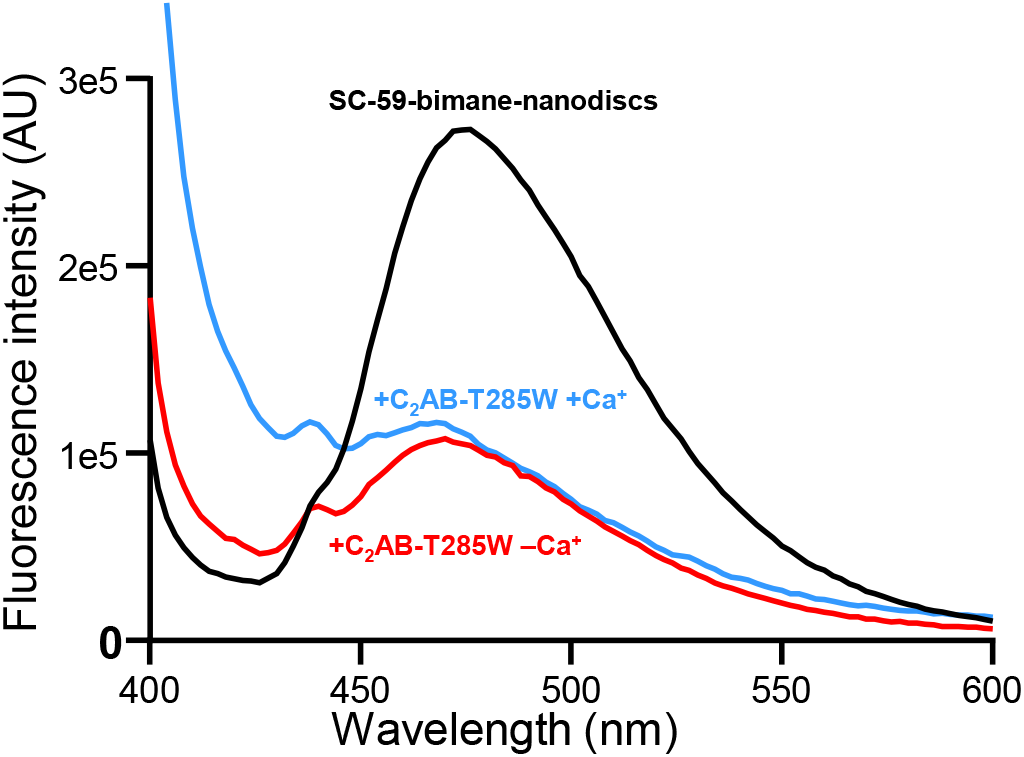
Ca^2+^ does not induce dissociation of Syt1 C_2_AB from membrane-anchored SNARE complex. The graph shows fluorescence spectra of nanodisc-anchored SC-59-bimane alone (black curve), or in the presence of C_2_AB-T285W and Mg-ATP plus 1 mM EGTA (red curve) or 1 mM Ca^2+^ (blue curve). Mg-ATP was included in the samples to hinder non-specific C_2_AB-SNARE complex interactions (47).

### Region II of the Syt1-SNARE complex remains bound upon disruption of region I

In parallel with our MD simulations and bimane fluorescence experiments, we studied the effects of mutations in the primary interface on Syt1 C_2_B domain-SNARE complex binding in solution using NMR spectroscopy to probe the energy landscape of the primary interface and to correlate binding through this interface with Syt1 function in the electrophysiological data (52). For this purpose, we acquired transverse relaxation optimized spectroscopy (TROSY)-enhanced ^1^H-^15^N heteronuclear single quantum coherence (HSQC) spectra of wild type (WT) and mutant versions of ^2^H,^15^N-labeled C_2_B domain specifically ^13^CH_3_- labeled at the Ile δ1 and Met methyl groups (here referred to as ^15^N-C_2_B for simplicity), and analyzed the cross-peak perturbations caused by titration with SNARE complex four-helix bundle bound to a complexin-1 fragment that spans residues 26-83 and prevents aggregation of C_2_B-SNARE complexes (referred to as CpxSC). Previous studies using this methodology showed that, in solution: i) the WT C_2_B domain binds to CpxSC via both the primary interface and the polybasic region that interacts with lipids when the SNARE complex is membrane-anchored; ii) an R322E/K325E mutation in the polybasic region (REKE) abolishes CpxSC binding through this region; iii) an R398Q/R399Q mutation in region II of the primary interface (RQRQ) abolishes binding through this interface; iv) an R322E/K325E/R398Q/R399Q mutation abolishes all binding; and v) an E295A/Y338W mutation in region I enhances the affinity of binding through the primary interface (47).

Superpositions of full ^1^H-^15^N TROSY-HSQC spectra of WT and mutant ^15^N-C_2_B titrated with CpxSC are shown in Fig. S6-S14. The positions of residues corresponding to diagnostic cross-peaks that reflect binding to region I or II of the primary interface are illustrated in red on the ribbon diagram of Fig. S15A, which displays the side chains of the C_2_B domain that were mutated in stick models. R281 is one of the three arginines in region II and was mutated to compare its contributions to C_2_B-CpxSC binding with those of R398 and R399; A402 is located between regions I and II; and E295 and Y338 are the key residues of C_2_B that form region I. These four residues were replaced individually in mutations that suppress the dominant negative lethality caused by overexpression of Syt1 with disrupted C_2_B domain Ca^2+^ binding sites in *Drosophila* (50). In initial experiments we analyzed the effects of a double E295K/Y338A mutation in the background of WT C_2_B domain and observed that the V283 and K288 cross-peaks still shifted upon titration with CpxSC, showing that binding to region II still persisted (Fig. S7, S15B). Since Y338 was replaced by aspartate in the mutant that suppressed lethality in the *Drosophila* screen, and an A402T mutation also suppressed lethality (50), we titrated a double Y338D/A204T ^15^N-C_2_B domain mutant but still observed binding to region II (Fig. S8, S15B). In these experiments it was difficult to assess the effects on binding to region I because of the limited cross-peak shifts for residues of this region observed for WT C_2_B (Fig. S6, S15B). The limited nature of these changes arises because of interference by binding modes involving the polybasic region and, consequently, full binding to the primary interface is better observed for the REKE mutant, which exhibits more overt shifts for many cross-peaks, including some from residues in region I (47) such as those of K297 and L299 (Fig. 4, S9; compare with WT C_2_B in Fig. S6, S15B). Hence, additional mutations in the primary interface were performed in the background of the REKE mutant.

**Figure 4.**
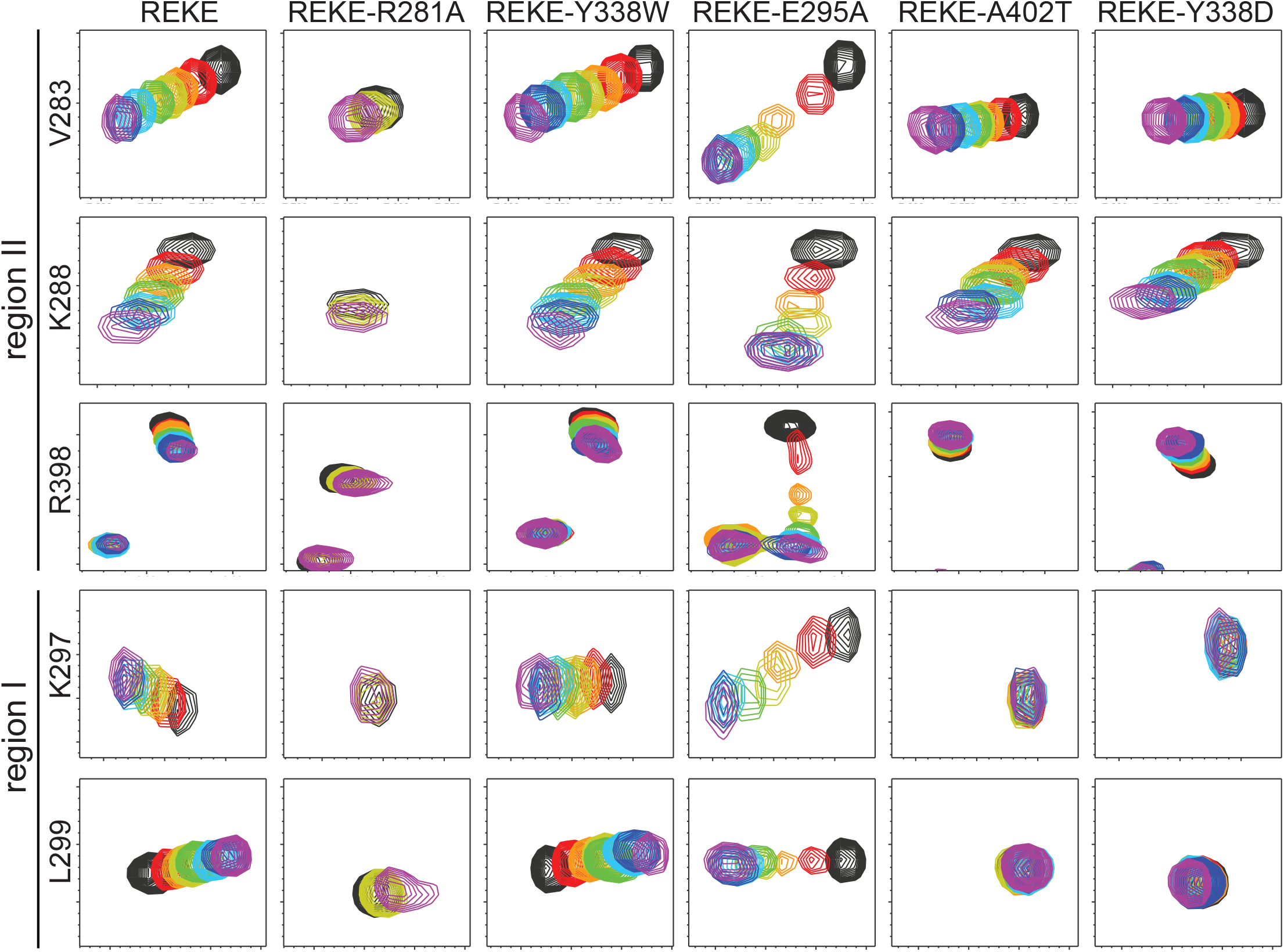
The Syt1 C_2_B domain still binds to the SNARE complex through region II of the primary interface when binding through region I is abolished. The diagrams show expansions of ^1^H-^15^N TROSY HSQC spectra of ^15^N-C_2_B domain mutants (as indicated above) acquired in isolation (black contours) or increasing concentrations of CpxSC (rainbow colours). The residues corresponding to the cross-peaks shown in the expansions and the regions where they are located are indicated on the left. The following concentrations of ^15^N-C_2_B mutant and CpxSC (μM/μM) were used for the different mutants (from black to purple): REKE 32/0, 30/10, 28/19, 26/28, 24/36, 20/51, 17/64, 12/85; REKE-Y338W 32/0, 30/10, 28/19, 26/28, 24/36, 20/53, 17/67, 12/88; REKE-E295A 32/0, 30/10, 28/19, 26/28, 24/36, 20/53, 17/65, 12/88; REKE-A402T 32/0, 30/10, 28/19, 26/28, 24/36, 20/53, 17/67, 12/88; REKE-Y338D 32/0, 30/10, 28/19, 26/28, 24/36, 20/53, 17/68, 12/88. For the REKE-R281A mutant the concentrations were 32/0 (black), 26/28 (yellow) and 13/98 (purple). The intensities of cross-peaks decreased as CpxSC was added because ^15^N-C_2_B was diluted and binding to CpxSC causes cross-peak broadening. Contour levels were adjusted to compensate for these decreased intensities, but after doing comparable adjustments for the spectra of the REKE-E295A mutant the cross-peaks in the middle of the titration are still very weak because of chemical exchange broadening.

Replacement of R281 with alanine (REKE-R281A mutant) strongly impaired overall binding of the C_2_B domain to CpxSC, although some binding remained at high CpxSC concentrations (Fig. 4, S10). These effects are comparable to those caused by individual R398Q or R399Q mutations but milder than those caused by the double R398Q/R399Q mutation (47). These results suggest that each arginine has comparable contributions to SNARE binding, consistent with the similar phenotypes caused by point mutations in the arginines (52). We also analyzed REKE-Y338W and REKE-E295A ^15^N-C_2_B mutants to dissect the contributions of each residue substitution to the enhancement in CpxSc binding affinity caused by the E295A/Y338W mutation in region I of the C_2_B domain (47). We found that the cross-peaks shifts observed for REKE-Y338W were similar to those observed for REKE (Fig. 4, S11), consistent with the lack of a phenotype caused by this mutation (52). In contrast, the E295A mutant exhibited more extensive shifts throughout the ^1^H-^15^N TROSY-HSQC spectrum than the REKE mutant Fig. S9, S12), indicating a more intimate interaction, and the shifts in diagnostic cross-peaks were larger earlier in the titration than observed for REKE (Fig. 4). We even observed clear chemical exchange broadening for some cross-peaks of the E295A mutant (e.g. the R398 cross-peak, Fig. 4) and broadening beyond detection of the W404 side chain cross-peak (Fig. S12). These observations show that the E295A mutation enhances the affinity of the C_2_B domain for Cpx-SC whereas the Y338W mutation has little effect. Indeed, the K_D_s between C_2_B and CpxSC estimated by fitting the changes in the ^1^H chemical shift of V283 and in the ^15^N chemical shift of K288 to a single binding site model (Fig. S16) were 34.4 ± 9.3 and 33.2 ± 4.6 μM for the REKE mutant, 30.4 ± 4.4 and 33.0 ± 8.5 μM for REKE-Y338W, and 7.3 ± 2.0 and 4.6 ± 0.4 μM for REKE-E295A. The about 5 to 6-fold enhancement in the affinity of REKE-E295A derived from these results correlates exquisitely well with the 4-fold decrease in spontaneous release caused by the E295A mutation (52), strongly supporting the notion that Syt1-SNARE binding through the primary interface inhibits spontaneous release. Moreover, the E295A and E295/Y338W mutations cause similar phenotypes, including enhanced vesicle priming and severe impairments in Ca^2+^-triggered release (52), showing that the phenotypes of the double mutant observed earlier (25) are caused by the E295A substitution.

No changes were caused by CpxSc on the K297 and L299 cross-peaks of ^15^N REKE-C_2_B bearing either A402T or Y338D mutations (REKE-A402T and REKE-Y338D) (Fig. 4, S13-S14), showing that the A402T and Y338D mutations practically abrogate binding to region I. Intriguingly, the changes in the V283 and K288 cross-peaks of both mutants were comparable to those observed for REKE-C_2_B, showing that robust binding to region II remained for these mutants, albeit the binding mode was likely altered given the different direction of the movements of the R398 cross-peak for REKE-A402T and REKE-Y338D compared to REKE (Fig. 4). Hence, binding through region II remains even in the absence of binding through region I.

### Analysis of interactions between C_2_B domain arginines and SNARE acidic residues at region II by MD simulations

The model postulating that Ca^2+^-induces dissociation of Syt1 from the SNARE complex to trigger neurotransmitter release arose in part because of the observation that the E295A/Y338W mutation in region I of the primary interface impairs release (25) but enhances C_2_B domain-SNARE complex binding (47). Our NMR results showing that disruption of region I does not lead to dissociation of region II, together with the fluorescence data showing that Ca^2+^ does not dissociate C_2_AB from liposome-anchored SNARE complex (Fig. 3), bring the possibility that neurotransmitter release is triggered by a Ca^2+^-induced re-arrangement of the primary interface in which region I dissociates but region II does not. To develop a hypothesis of how such re-arrangement could trigger release, it is important to have a clear picture of the configuration of the C_2_B domain with respect to the SNARE complex and the membrane. Such a model could be derived from the multiple crystal structures of Syt1-SNARE complexes that have been described (25, 49), which revealed similar configurations of the primary interface. There was some variability in the interactions involving the three C_2_B arginines of region II (R281, R398, R399) in the various structures, but the arginines were generally close to multiple acidic residues of both SNAP-25 and syntaxin-1 (Table S2) (Fig. S17A,B). However, MD simulations of primed Syt1 C_2_AB-SNARE-complexin-1 complexes bridging a vesicle and a flat bilayer suggested that additional configurations exist in which the three arginines interact primarily with acidic residues of SNAP-25, perhaps favored by simultaneous interactions of the C_2_B domain with the flat bilayer (27). A concern about these simulations is their limited length and the fact that the SNARE four-helix bundles were in distinct stages of assembly, which could affect the interactions with Syt1.

To further examine the consistency of these results, we set up a system with four almost fully assembled SNAREs complexes bridging a vesicle and a flat bilayer, and bound to Syt1 C_2_AB through the primary interface as in the crystal structure corresponding to PDB code 5KJ7 (Fig. S18A). Complexin-1 was omitted to examine whether the previously observed Syt1-SNARE binding modes (27) are altered in its absence. We ran a 596 ns simulation (referred to as s1action2), which led to similar configurations of the C_2_B domain with respect to the SNARE complex and the flat bilayer (Fig. S18B-E) that resembled those observed in the previous simulations (27). This conclusion was supported by analysis of the distances between the guanidine carbon (CZ) of R281, R398 and R399 of the four C_2_AB molecules and the carboxyl carbon of the nearby acidic residues (CG for aspartates, CD for glutamates) of syntaxin-1 or SNAP-25 along the trajectories (Fig. S19). These plots and the trajectory-averaged distances calculated for each C_2_AB-SNARE complex (Table S3) again showed a predominance of configurations in which the three arginines interact primarily with the SNAP-25 acidic residues, as illustrated by a representative pose shown in Fig. S17C,D), although there were also frequent interactions with syntaxin-1 acidic residues. The similarity of the distances averaged over the four C_2_AB-SNARE complexes in the s1action2 simulation and those calculated for the prsg simulation of ref. (27) (Table S3) further shows the consistency of these results and suggest that the Syt1-SNARE binding modes are not substantially affected by complexin-1.

### A lever hypothesis for Syt1 action

How can Ca^2+^ binding to the C_2_B domain play such a critical role in triggering fast neurotransmitter release (33, 34) if its Ca^2+^-binding loops are pointing away from the site of fusion in the primed state? The hypothesis that we propose to answer this question emerged from the realization that zippering of the jxt linkers separating the SNARE motifs and TM regions of synaptobrevin and syntaxin-1 is critical for release (63, 64) and liposome fusion (65), which was reinforced by our recent MD simulations (15). The simulations showed that, because the SNARE four-helix bundle is oriented parallel to the bilayers and the TM regions are approximately perpendicular, the jxt linkers must form kinked helices (e.g. Fig. 5A) or become partially unstructured (e.g. Fig. S18B-E) to accommodate the changes in direction. Linker zippering led to fast (microsecond scale) fusion in the simulations (15) and is in principle energetically favorable (66). However, linker zippering is expected to be hindered by substantial energy barriers because the zippering pulls the hydrophobic TM regions into the polar bilayer interface (67), and the abundant basic and aromatic residues of the jxt linkers have high propensities to interact with the lipids (15, 27). Our hypothesis predicts that Ca^2+^ binding to the Syt1 C_2_B domain helps to overcome these energy barriers because it induces reorientation of the domain with respect to the flat membrane, from the parallel orientation existing in the primed state (modeled in Fig. 5A) to an approximately perpendicular orientation that allows insertion of both of its Ca^2+^ binding loops into the membrane as observed by EPR (54) (modeled in Fig. 5B). During this reorientation, the C_2_B domain is predicted to act as a lever, as region I of the primary interface dissociates but the three arginines of region II remain electrostatically bound to the SNARE acidic residues and pull the SNARE four-helix bundle away from the fusion site, facilitating linker zippering (Fig. 5B) and fast membrane fusion. This model explains how Syt1 can act remotely from the site of fusion without inserting its Ca^2+^-binding loops near this site, where Syt1 would hinder the action of the SNAREs that trigger membrane fusion. The model also allows rationalization of the various phenotypes caused by mutations in the primary interface, most notably the findings that neurotransmitter release is impaired by mutations that impair or enhance binding through this interface and is differentially affected by mutations in the SNARE acidic residues (25, 52, 53, 68) (see discussion).

**Figure 5.**
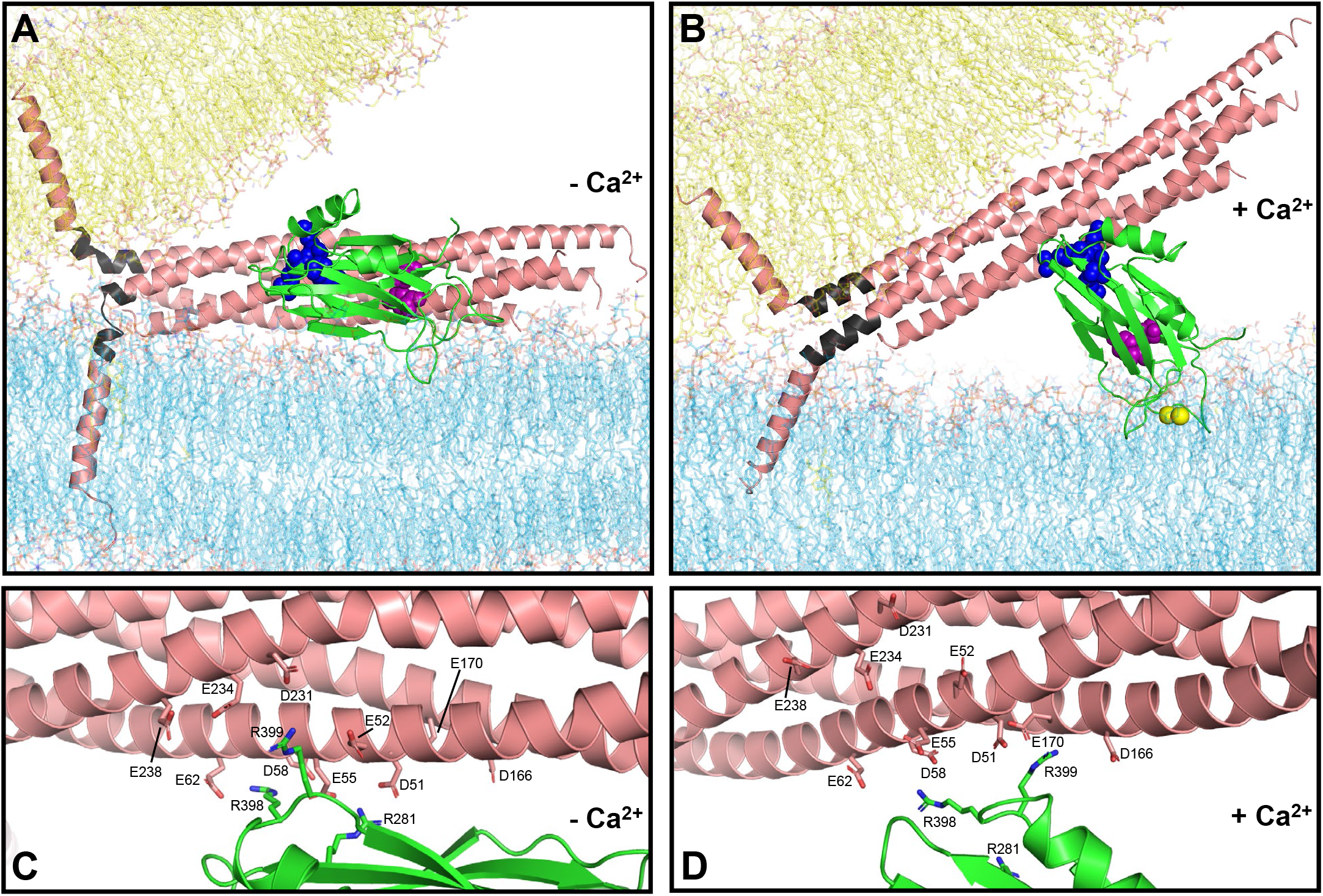
Lever hypothesis of Syt1 function. (*A*) Model of the primed state before Ca^2+^ influx. Lipids are shown as stick models with the same color coding as in Fig. 1. The SNARE complex is represented by a ribbon diagram in salmon color except for the jxt linkers, which are colored in dark gray. The Syt1 C_2_B domain is represented by a green ribbon diagram with key residues at the primary interface shown as spheres: E295 and Y338 (region I) are in magenta; R281, R398 and R399 (region II) are in blue. The SNARE complex, flat bilayer and vesicle configurations were extracted from one of the frames of the fusion2g simulation described in (15) to illustrate a potential configuration of the primed state. The C_2_B domain was placed at a location analogous to that in one of the primed complexes at the end of the s1action2 simulation. The Syt1 C_2_A domain and complexin are also part of the primed state but are not shown for simplicity. (*B*) Model of a potential Ca^2+^-activated state built manually in Pymol using the configuration of panel (*A*) as starting point, rotating the C_2_B domain to an approximately perpendicular orientation with respect to the flat bilayer that mimics the plasma membrane, as observed by EPR (54), and moving the SNARE four-helix bundle to keep ionic interactions of the SNARE acidic residues with the arginines of region II of the C_2_B domain. The synaptobrevin and syntaxin-1 jxt linkers were modeled as helices that extend from the four-helix bundle based on the crystal structure of the SNARE complex (77), and the TM regions were pulled toward the polar membrane-membrane interface to maintain the continuity of the polypeptide chains. Ca^2+^ ions are shown as yellow spheres. (*C, D*) Close-up views of the primary interface in panel (*A*) and (*B*), respectively, showing the C_2_B arginines and SNARE acidic residues as stick models to show the interactions between these residues in the proposed initial configuration and in one of many potential configurations of the Ca^2+^-activated state.

### Ca^2+^ -induces reorientation of the Syt1 C_2_B domain bound to membrane-anchored SNARE complex

Testing our hypothesis is hindered by several factors. First, the proposed Ca^2+^-induced reorientation of the C_2_B domain may occur transiently to trigger release in microseconds and hence may be difficult to observe with experiments performed under equilibrium conditions. Second, the reorientation may be difficult to monitor because it may be much less pronounced than depicted in Fig. 5 (see discussion). Third, FRET studies showed that Syt1 C_2_AB interacts with nanodisc-anchored SNARE complex in part through the primary interface, but the binding is weak and there are also non-specific binding modes that muddle the interpretation of the data (47). Buffers with physiological ionic strength and including ATP hinder such unwanted interactions but also decrease the overall affinity of Ca^2+^-free C_2_AB for nanodisc-anchored SNARE complex. Ca^2+^ enhances the overall affinity, but the enhancement arises because Ca^2+^ strongly increases the affinity of C_2_AB for the lipids. Interactions between C_2_AB and SNARE complex anchored on nanodiscs are actually weakened by Ca^2+^, as shown by the fact that C_2_AB binds with similar affinity to nanodiscs containing or lacking SNARE complex (47). Fourth, use of bimane fluorescence quenching by a nearby tryptophan (Fig. 3) to monitor the postulated Ca^2+^-induced reorientation is hindered by the need to place the tryptophan near the bimane group, which might alter the native binding mode and potentially enhance affinity due to tryoptophan-bimane interactions.

Taking these concerns into account, we studied whether Ca^2+^ induces reorientation of the C_2_B domain bound to liposome-anchored SNARE complex using FRET experiments with Syt1 C_2_AB covalently linked through a 37-residue sequence to the C-terminal SNARE motif of SNAP-25, an approach that was used to stabilize Syt1-SNARE complex interactions in the X-ray studies that revealed the primary interface (25). The covalent link facilitates quantitative binding of C_2_AB to the SNARE complex through the primary interface in the absence of Ca^2+^ and thus prevents FRET changes that could arise from a Ca^2+^-induced increase of the overall affinity of C_2_AB for the SNARE complex-containing liposomes rather than from C_2_B reorientation. We also include complexin-1 in these experiments because it hinders non-specific interactions between Syt1 and the SNARE complex (47). A photostable Cy5-Trolox acceptor probe was attached to residue 412 of C2AB in the fusion protein, which was used to form liposome-anchored SNARE complexes that were also labeled with a Cy3-Trolox donor probe on residue 27 or 34 of the SNAP-25 N-terminal SNARE motif. These positions were chosen because the distances between donor and acceptor probes are predicted to change substantially based on the model of Fig. 5 (see also Fig. 6A). To have reference spectra without FRET, we acquired fluorescence spectra with analogous samples in the presence of detergent to disrupt the liposomes and of NSF plus αSNAP to disassemble the SNARE complex.

**Figure 6.**
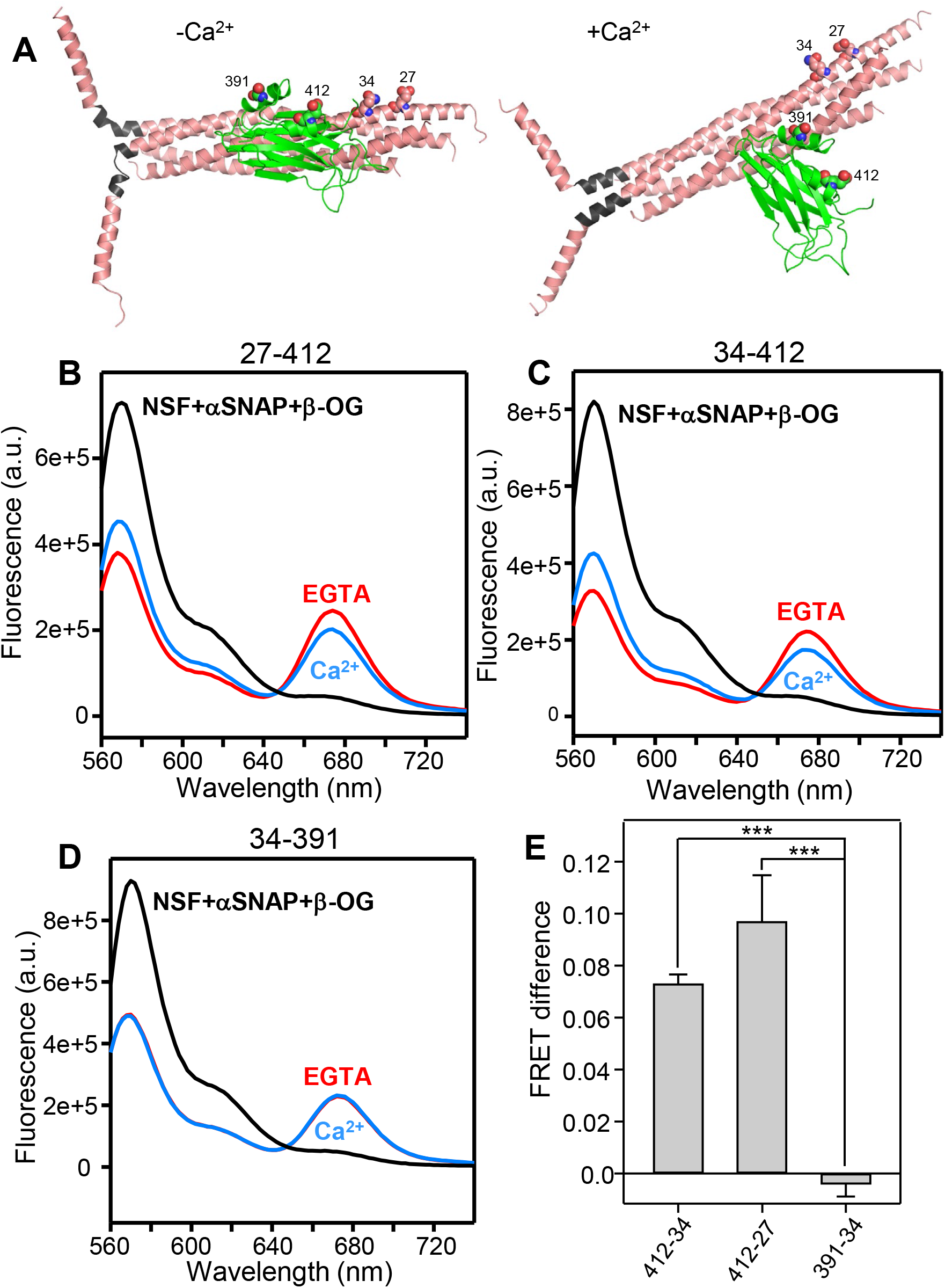
Ca^2+^ induces reorientation of the Syt1 C_2_B domain with respect to membrane-anchored SNARE complex. (*A*) Models of the Syt1 C_2_B domain-SNARE complex in the primed state before Ca^2+^ influx and in a potential Ca^2+^-activated state, analogous to those of Fig. 5A,B and with the same color code. Residues of C_2_B and SNAP-25 that were mutated to cysteines to attach fluorescent probes are shown as spheres and labeled with the residue number. In these models, the following changes are predicted for the distances between the approximate points of attachment of the fluorescent probes in the absence or presence of Ca^2+^: SNAP-25 Q34 CD carbon - Syt1 E412 CD carbon from 34 Å (-Ca^2+^) to 48 Å (+Ca^2+^); SNAP-25 E27 CD carbon - Syt1 E412 CD carbon from 41 Å (-Ca^2+^) to 52 Å (+Ca^2+^); SNAP-25 Q34 CD carbon - Syt1 E391 CB carbon 31 Å (-Ca^2+^) to 28 Å (+Ca^2+^). Note that there is a large uncertainty in the distance changes because the configuration of the Ca^2+^-activated state is unknown and the 27-412 and 34-412 distances depend strongly on the angle of rotation (Supplementary Discussion). (*B-D*) Fluorescence spectra of liposome-anchored C_2_AB-SNARE-complexin-1 complex with stabilized Cy3 and Cy5 probes attached to the following residues of SNAP-25 and the C_2_B domain, respectively: 27-412 (*B*), 34-412 (*C*) and 34-391 (*D*). The spectra were obtained in the presence of Mg-ATP plus 1 mM EGTA (red curves), 1 mM Ca^2+^ (blue curves) or 1% β-OG plus NSF and αSNAP as a control to disrupt the Syt1-SNARE complex (black curves). In the experiments of (*B-D*), Mg-ATP was included in the samples to hinder non-specific C_2_AB-SNARE complex interactions (47). (*E*) Quantification of the FRET difference between the spectra obtained in 1 mM Ca^2+^ and 1 mM EGTA normalized with the FRET observed in the presence of 1 mM EGTA. Bars represent mean values and error bars represent standard deviations from experiments performed at least in triplicate. Statistical significance and p values were determined by one-way analysis of variance (ANOVA) with the Holm-Sidak test (*** p < 0.001).

Importantly, comparison of the spectra obtained in EGTA with the reference spectra revealed robust FRET for the samples of liposome-anchored C_2_AB-SNARE-complexin-1 complex with the 27-412 and 34-412 FRET pairs and Ca^2+^ induced reproducible decreases in FRET (Fig. 6B,C), which was verified with experiments performed on different days with different samples. As a negative control, we also performed FRET experiments with Cy5-Trolox attached to residue 391 of C_2_B in the fusion protein and Cy3-Trolox attached to residue 34 of SNAP-25, since the distance between the two probes was predicted to change little upon Ca^2+^ binding (Fig. 6A). Indeed, we did not observe appreciable changes in FRET for the 34-391 pair (Fig. 6D), supporting the prediction and confirming that Ca^2+^ does not induce dissociation of Syt1 from the liposome-anchored SNARE-complexin-1 complex under these conditions. Quantification of the Ca^2+^-induced changes in donor fluorescence for repeat experiments showed the statistical significance of these results (Fig. 6E). We note that the changes in distances between the fluorescent probes predicted by the model are approximate given the considerable size of the probes and the uncertainty in the configuration of the Ca^2+^-bound state (Supplementary Discussion). Regardless of this uncertainty, these FRET data clearly support the notion that Ca^2+^ induces reorientation of the C_2_B domain bound to membrane-anchored SNARE complex, which is at the heart of the model of Syt1 action that we propose.

## Discussion

The function of Syt1 as the Ca^2+^ sensor that triggers neurotransmitter release is well established, but its mechanism of action has remained enigmatic despite intense research for over three decades. Models postulating that Syt1 cooperates with the SNAREs in triggering membrane fusion by perturbing bilayers, inducing membrane curvature or bridging the two membranes were attractive, but appeared to be incongruous with the finding that Syt1 binds to the SNARE complex through the primary interface (25), which orients the C_2_B domain Ca^2+^-binding loops away from the fusion site. The proposal that Ca^2+^ induces dissociation of Syt1 from the SNARE complex, which would allow the C_2_ domains to reorient toward the site of fusion, provided a potential solution to this paradox (47) but did not bode well for the fast speed of release. Moreover, it was unclear how these models of Syt1 function can be reconciled with our MD simulation revealing microsecond scale fusion induced by SNARE complexes upon jxt linker zippering (15). The MD simulations presented here now cast further doubt on these models and suggest that Syt1 does not act directly at the site of fusion. Our NMR titrations, together with previous EPR data (54), lead naturally to the hypothesis that Ca^2+^ binding induces reorientation of the C_2_B domain on the membrane and partial rather than full dissociation from the SNAREs such that ionic interactions involving the C_2_B arginine cluster pull the SNARE complex and facilitate jxt linker zippering to induce fast membrane fusion. Although further research will be required to test this lever hypothesis and the underlying details, the notion that Ca^2+^-induced remodeling of Syt1-SNARE interactions at the primary interface is crucial for neurotransmitter release is supported by our FRET assays and by the correlation of our MD simulation and NMR results with the electrophysiological data described in the accompanying paper (52).

### Need to revise current models of Syt1 action

MD simulations need to be interpreted with caution because force fields are not perfect and because of the limited simulation lengths that can be achieved, particularly for multimillion atom systems such as those studied here. However, all-atom MD simulations have provided a powerful tool to visualize the neurotransmitter release machinery (27, 56) and to elucidate how the SNAREs triggers fast membrane fusion, showing how linker zippering pulls the hydrophobic TM region to the polar membrane-membrane interface where they catalyze lipid acyl chain encounters at the interface, initiating bilayer merger (15). This mechanism makes a lot of sense from a physicochemical perspective and explains a large amount of experimental data (15) but does not involve a central role for membrane curvature in initiating fusion, in contrast to assumptions made from theoretical calculations (69). Hence, there is no need for neurotransmitter release to depend on induction of membrane curvature by Syt1, as proposed by some models (39, 40). Other models postulated that insertion of the Syt1 C_2_ domains perturbs bilayers and causes lipid disorder (35, 37, 38), which can facilitate fusion, but bilayer perturbation is efficiently caused by the events that ensue after SNARE jxt linker zippering without the need for Syt1 (15). In fact, our MD simulations strongly suggest that, when placed near the site of fusion, the Syt1 C_2_ domains hinder SNARE action [Fig. 1, S1-S5; see also (27)]. Note also that the experiments supporting these models of Syt1 function used much larger protein-to-lipid ratios than the Syt-to-lipid ratios present in synaptic vesicles (57), and our MD simulations indicate that a few Syt1 molecules induce rather limited perturbation of lipid bilayers (e.g. Fig. 2).

An additional model predicted that the Syt1 C_2_ domains bridge the two membranes, helping the SNAREs to bring the membranes together (42, 43). This notion was inspired in part by the observation that Syt1 C_2_AB can indeed bridge two membranes and in part by the overall idea that substantial energy is required to bring two membranes together (42, 70). However, the extended membrane-membrane interfaces induced by SNARE complexes, even when full zippering is prevented (60), and our previous MD simulations (15, 27), indicate that bringing membranes into contact is easier than previously thought. Moreover, the Syt1 C_2_ domains clearly hindered the ability of the SNAREs to bring the membranes together in our sytfusion3 simulation (Fig. 1D, S5). In summary, previous models of Syt1 function were attractive but are not supported by the new perspective on membrane fusion emerging from the MD simulations. Moreover, the fact that binding of the Syt1 C_2_B domain to the SNARE complex through the primary interface places its Ca^2+^-binding loops away from the site of fusion argues strongly against these models.

### Multiple roles of the primary interface

Previous studies (25, 50, 53), together with the correlations between our binding data (47) (Fig. 4, S6-S16) and electrophysiological results (52), provide overwhelming evidence that the primary interface plays central roles in vesicle priming, clamping of spontaneous release and Ca^2+^ triggering of release. The strong disruption of vesicle priming and evoked release caused by the R398Q/R399Q mutation (25, 52, 53) that abolishes binding through region II of the primary interface (47) shows the critical functional importance of this region. The Y338D and A402T mutations that abolish C_2_B-SNARE binding through region I (Fig. 4, S13, S14) also impair priming and evoked release (52), but the E295A and E295A/Y338W mutations in region I that enhance overall binding (47) (Fig. 4, S12) strongly impair evoked release while enhancing priming (25, 52). These findings show that interactions involving region I mediate priming but need to be dissociated for Ca^2+^ triggering of release. The primary interface is also important to inhibit spontaneous release, as mutations in this interface that impair binding generally lead to enhanced spontaneous release, and the E295A mutation causes a 5-6-fold enhancement in binding (Fig. 4, S16) and a 4-fold decrease in spontaneous release (52). The finding that the E295A substitution mediates the enhanced binding and the phenotypes caused by the double E295A/Y338W mutation (52) (Fig. S4, S11, S12) might seem surprising because E295 forms a salt bridge in crystal structures of Syt1-SNARE complexes (25, 49), but can be attributed to the increase in the positive electrostatic potential of C_2_B caused by the mutation, as overall electrostatics play a key role in binding of the highly basic C_2_B domain to the highly acidic SNARE complex (71).

### The lever hypothesis

The proposal that Ca^2+^ induces dissociation of Syt1 from the SNARE complex arose in part because the perpendicular orientation of the C_2_B domain with respect to the membrane induced by Ca^2+^ (54) is incompatible with binding of C_2_B to the SNARE complex via the full primary interface, and in part to explain the impairment of release caused by the E295A/Y338W mutation despite enhancing binding (47). The observation that robust C_2_B-SNARE complex binding through region II of the primary interface remains even upon disruption of region I with the Y338D mutation (Fig. 4, S14) provided a key clue suggesting that Ca^2+^ induces partial rather than full dissociation of the primary interface. Thus, reorientation of C_2_B with respect to the membrane can occur while interactions mediated by region I are released but those involving region II remain. In support of this notion, our bimane fluorescence quenching results (Fig. 3) and the FRET data obtained with the 34-391 pair (Fig. 6D) clearly show that Ca^2+^ does not dissociate the C_2_AB fragment from membrane-anchored SNARE complex, and the FRET data obtained with the 27-412 and 34-412 pairs (Fig. 6B,C) show that Ca^2+^ indeed alters the orientation of the C_2_B domain with respect to the SNARE complex. The exact nature and extent of the reorientation is still unclear (Supplementary discussion) but, since only one of the two Ca^2+^-binding loops is inserted into the membrane in the absence of Ca^2+^ (Fig. 5A), the reorientation is most likely driven by insertion of the other Ca^2+^-binding loop into the membrane (Fig. 5B) together with Ca^2+^-phospholipid interactions that complete the coordination spheres of the Ca^2+^ ions (36).

Ca^2+^-induced reorientation of the C_2_B domain is expected to pull the SNARE complex away from the fusion site through electrostatic interactions of the three C_2_B arginines of region II with acidic residues of the SNARE complex. The importance of these interactions for Ca^2+^ triggering of release is supported by the fact that individual R281A, R398Q and R399Q mutations in Syt1 and D51N and E55Q mutations in SNAP-25 severely impair evoked release without affecting vesicle priming, thus lowering the vesicle release probability (52). Intriguingly, mutations in E52 of SNAP-25 or acidic residues of syntaxin-1 (E228, D231, E234 and E238) increase the release probability (52, 72). These distinct phenotypes can be rationalized by the observation that region II of the primary interface has two faces. D51 and E55 are located along one side of the SNAP-25 N-terminal SNARE motif helix, whereas SNAP-25 E52 and the syntaxin-1 acidic residues are located on the other side of the helix (Fig. S17A,C). In the configurations most populated during our MD simulations of the primed state, the C_2_B arginines are generally closer to the former face, and R281 interacts with D51 and E55 while R398 interacts with E55, D58 and E62 (Fig. S17C,D, S19, Table S3). The observed phenotypes indicate that this is the active face involved in the pulling action of Syt1 that triggers evoked release. Configurations with C_2_B arginines closer to the syntaxin-1 acidic residues on the other face of region II also occur in our simulations (Fig. S19) and are observed in crystal structures (Fig. S17A,B, Table S2), but manual inspection suggests that the syntaxin-1 helix needs to move away from the C_2_B domain arginines as C_2_B reorients and pulls the SNARE complex (Fig. 5D). Hence, this face of region II may hinder activation of evoked release by Syt1, which would explain the increased release probabilities caused by mutations in SNAP-25 E52 or acidic residues of syntaxin-1. We note that R399 is oriented toward this inhibitory face (Fig. S17A,C) and yet is important for evoked release (52). Since interactions of the arginines must be remodeled during reorientation of the C_2_B domain, it is plausible that remodeling leads to new interactions of R399 that are key for the pulling action that triggers release. For instance, modeling suggests that R399 may interact with D166 and E170 in the active state (Fig. 5D), which might explain the strong impairments in neurotransmitter release caused by mutations in these residues (68).

Pulling the SNARE complex upon reorientation of C_2_B should increase the angle between the long axis of the four-helix bundle and the plasma membrane, consistent with data showing that the angle between the SNARE complex and a support lipid bilayer is increased by binding to Ca^2+^-saturated Syt1 C_2_AB (73). Such tilting may enable formation of a continuous helix in syntaxin-1 (Fig. 5B), which might facilitate fusion. However, it is most likely that the key consequence of the C_2_B domain reorientation is that pulling the SNARE complex away from the site of fusion facilitates extension of the syntaxin-1 and synaptobrevin SNARE motif helices into the jxt linkers, which is critical for neurotransmitter release (63, 64) and liposome fusion (65). This notion emerged in part from MD simulations showing that jxt linker zippering can mediate fast, microsecond scale membrane fusion, but zippering is hindered by the natural geometry of the system, which dictates that the directions of the polypeptide chains turn the corner at the jxt linkers (15) (Fig. 5A). It is unclear whether the jxt linkers form kinked helices in the primed state, as in Fig. 5A, or adopt more disordered structures before Ca^2+^ influx, but there is little doubt that the abundant basic and aromatic residues in the linkers interact with the lipids in the primed state (27) and that such interactions need to be rearranged for the linkers to zipper. The central aspect of our model is that the pulling force of the Syt1 C_2_B domain on the SNARE complex lowers the energy barrier for such rearrangement, facilitating linker zippering. This model naturally explains why the C_2_B domain Ca^2+^ binding loops are pointing away from the site of fusion when Syt1 binds to the SNARE complex through the primary interface, helping to trigger fast fusion remotely instead of being close to the fusion site where the C_2_ domains would hinder SNARE action (Fig. 1, S1, S2, S4, S5). We note that Ca^2+^ binding was proposed to induce a movement ‘en bloc’ of the Syt1-SNARE complex that deforms the membranes (25). While this model is inconsistent with the effects of mutations in region I of the primary interface on C_2_B-SNARE binding (Fig. 4) and neurotransmitter release (25, 52), the seminal crystal structure described in this study was crucial to develop the model proposed here.

Clearly, multiple aspects of our model are speculative and further research will be required to examine these aspects in detail and test the overall lever hypothesis of Syt1 function. For instance, the nature of the C_2_B domain reorientation is unclear and it might be transient or much less pronounced than predicted in Fig. 5, as small structural changes may be sufficient to overcome the energy barrier that hinders Ca^2+^-evoked release (Supplementary Discussion). An additional question is what is the role of the Syt1 C_2_A domain, as Ca^2+^ binding to this domain contributes to trigger neurotransmitter release even if it is not as critical as Ca^2+^ binding to the C_2_B domain (28, 32, 34). In three of the complexes of our MD simulations of the primed state, the C_2_A domain Ca^2+^ binding loops are next to those of the C_2_B domain, poised to bind concomitantly to the membrane upon Ca^2+^ binding (Fig. S18C-E), and in two of the complexes there are ionic interactions between acidic residues of SNAP-25 and a polybasic region of the C_2_A domain (K189-K192) (e.g. Fig. S20). Hence, the C_2_A domain could cooperate with the C_2_B domain in pulling the SNARE complex upon Ca^2+^-dependent membrane binding, and could provide a second anchor point on the membrane to exert force without sliding on the membrane surface, but these ideas need to be tested. Note also that other interactions of Syt1 have described, such as binding of the C_2_B domain to the SNARE complex through a so-called tripartite interface (49) or Ca^2+^-independent oligomerization (74). Although we and others have been unable to observe the tripartite interface in solution (47, 75) or the oligomerization by cryo-EM (7, 23, 76), the relevance of these interactions and their compatibility with our model need further investigation. The lever hypothesis and the underlying ideas presented here provide a framework to address these questions and elucidate how Syt1 triggers neurotransmitter release.

## Methods

Molecular dynamics simulations were performed using the same methodology employed in our previous simulations (15, 27). Methods used for site directed mutagenesis, expression and purification of SNARE proteins and Syt1 fragments, NMR spectroscopy, protein labeling with fluorescent probes and fluorescence spectroscopy were basically as described (47, 75). Specific details for all these methods are described in the *SI Appendix*.

## Acknowledgements

We thank Thomas Südhof amd Axel Brunger for extensive discussions. Most of the molecular dynamics simulations presented in this paper were performed through Pathways and Leadership Computing Resource allocations on Frontera at the Texas Advanced Computing Center of The University of Texas at Austin (URL: http://www.tacc.utexas.edu) (projects MCB20033 and IBN23002). This research also used computational resources provided by the BioHPC supercomputing facility located in the Lyda Hill Department of Bioinformatics, UT Southwestern Medical Center, TX (URL: https://portal.biohpc.swmed.edu). This work was supported by grant I-1304 from the Welch Foundation (to JR), by NIH Research Project Award R35 NS097333 (to JR), and by project 278001972-TRR186 from the Deutsche Forschungsgemeinschaft (DFG; German Research Foundation) (to CR).

## Data availability

All files corresponding to the experimental data presented in this paper and most files corresponding to the molecular dynamics simulations are being deposited in the dryad database. Because of the very large size of trajectory files, it was not practical to deposit them in this database, but these files are available from the corresponding author upon request.

## Supporting Information

### Supplementary Discussion

The model of Fig. 5A assumes that, in the primed state existing before Ca^2+^ influx, the Syt1 C_2_B domain binds to the SNARE complex through the primary interface and to the plasma membrane through the polybasic region, resulting in an approximately parallel orientation of the C_2_B domain with respect to the plasma membrane. The Ca^2+^-activated state of Fig. 5B was built manually in Pymol assuming that Ca^2+^-binding induces an approximately perpendicular orientation of the C_2_B domain with respect to the membrane, similar to that determined by electron paramagnetic resonance when C_2_B domain binds to SNARE-free membranes in the presence of Ca^2+^ (1), and that the reorientation pulls the SNARE complex through electrostatic interactions of the C_2_B domain arginines of region II with SNARE acidic residues. The changes in FRET observed for the 27-412 and 34-412 pairs, as well as the lack of FRET changes for the 34-391 pair (Fig. 6) show that Ca^2+^ indeed induces reorientation of the C_2_B domain with respect to liposome-anchored SNARE complex and agree qualitatively with the models of Fig. 5A,B. However, we did not attempt to do a quantitative comparison of the FRET changes with the distances between the probes predicted by the models (see legend of Fig. 6) because there is a large uncertainty in the configuration of the Ca^2+^-activated state and relatively small changes in the predicted angle of rotation of the C_2_B domain induced by Ca^2+^ lead to considerably large changes in the expected distances between the probes for the 27-412 and 34-412 pairs (but much smaller changes for the 34-391 pair). Moreover, the rotation might be transient or considerably smaller than that depicted in Fig. 5A,B, as small structural changes may be sufficient to trigger neurotransmitter release based on the energy considerations discussed below. In addition, a subset of the C_2_AB-SNARE-complexes used for the FRET measurements are located inside the liposomes and hence are inaccessible to the added Ca^2+^.

It is also worth noting that Syt1 C_2_AB binds with similar affinities to nanodiscs containing or lacking SNARE complex in the presence of Ca^2+^ (2), which suggested that C_2_AB does not interact with the membrane-anchored SNARE complex or that binding is rather weak. While our FRET data now show that C_2_AB does remain bound to membrane-anchored SNARE complex upon Ca^2+^ binding (Fig. 6D), a potential concern about our model is whether the underlying Syt1-SNARE interactions are too weak to trigger neurotransmitter release. In this context, it is important to consider that Ca^2+^ increases the rate of release in hippocampal neurons by about 18,000/70,000-fold (3, 4), which translates to a decrease in the energy barrier for release of 9.8-11.2 k_B_T. If at least three Syt1-SNARE complexes trigger release (5), the contribution of each complex to lower the energy barrier might be 3.3-3.7 k_B_T or less, which corresponds to a very weak binding energy and can be readily provided by electrostatic interactions and hydrogen bonds of the C_2_B domain arginines with the SNARE acidic residues in region II of the primary interface.

### Supplementary Methods

#### MD simulations

All-atom MD simulations were performed using Gromacs (6, 7) with the CHARMM36 force field (8). Solvation and ion addition for system setup were performed at the BioHPC supercomputing facility of UT Southwestern. Minimizations, equilibration steps and production molecular dynamics (MD) simulations were carried out on Frontera at the Texas Advanced Computing Center (TACC). Pymol (Schrödinger, LLC) was used for system design, manual manipulation and system visualization.

The methodology used to set up the systems and run MD simulations was analogous to that described previously (9, 10). The lipid compositions of the vesicle and the flat bilayers resembled those of synaptic vesicles and plasma membranes, respectively (11, 12), and are listed in Table S1 together with other parameters of the simulations. All systems were solvated with explicit water molecules (TIP3P model), adding potassium and chloride ions to reach a concentration of 145 mM and make the system neutral. All Syt1 C_2_AB molecules had five Ca^2+^ ions placed at the corresponding binding sites (13, 14) except for those of the s1action2 simulation, which was Ca^2+^-free. No restraints were used to keep the Ca^2+^ ions bound and no dissociation was observed during the simulations.

For all systems, we used the same vesicle generated previously (24) and moved lipids manually to accommodate different positions of the SNARE TM regions. The cac2absc simulation used the same bilayer and SNARE complexes used for the prsg simulation of (10). The four C_2_AB molecules were placed manually at positions interspersed between the four SNARE complexes, with the Ca^2+^-binding loops pointing toward the flat bilayer but without being in contact with it (Fig. S1A). The fusiong system was analogous to the cac2absc system but using four copies of the SNARE complex that was almost fully assembled in fusiong and placing the C_2_AB molecules with the Ca^2+^ binding loops pointing toward the center of the vesicle-flat bilayer interface (Fig. S2A). The nosytfusion system had the same initial configuration of the fusiong system but without C_2_AB molecules (Fig. S3A). The sytfusion2g system also had a configuration analogous to fusiong but with only two C_2_AB molecules that had the Ca^2+^-binding loops pointing in the same direction, toward the center of the vesicle-flat bilayer interface (Fig. S4A). The sytfusion3 system included a vesicle, a flat bilayer and four trans-SNARE complexes placed closer to the center of the bilayer-bilayer interface, compared to the systems of Fig. S1A-S4A, as in the fusion2g simulation of (9). A restrained MD simulation was used to pull the syntaxin-1 TM regions to designed positions such that the distance between the vesicle and the flat membrane was increased by 1.6 nm to allow sufficient space to place four C_2_AB molecules poised to bridge the two membranes (Fig. S5A). The s1action2 system was built with a vesicle and the same flat bilayer used for the prs2 simulation of (10), which is larger than that of the other simulations described here to provide sufficient space for interaction with C_2_AB molecules. The four trans-SNARE complexes were almost fully assembled and were placed at a distance from the bilayer-bilayer interface that was intermediate between those used for the prs2 simulation of (10) and for the fusion2g simulation of (9). Each SNARE complex had a C_2_AB bound through the primary interface (Fig. S18A).

Systems were energy minimized using double precision, whereas the default mixed precision was used in all MD simulations. The systems were heated to the desired temperature running a 1 ns simulation in the NVT ensemble with 1 fs steps, and then equilibrated to 1 atm for 1 ns in the NPT ensemble with isotropic Parrinello-Rahman pressure coupling (15) and 2 fs steps. NPT production MD simulations were performed for the times indicated in Table S1 using 2 fs steps, isotropic Parrinello-Rahman pressure coupling and a 1.1 nm cutoff for non-bonding interactions. Three different groups of atoms were used for Nose-Hoover temperature coupling (16): i) protein atoms; ii) lipid atoms; and iii) water and KCL ions. Periodic boundary conditions were imposed with Particle Mesh Ewald (PME) (17) summation for long-range electrostatics.

#### Protein expression and purification

The rat synaptobrevin-2 SNARE motif (residues 29–93), rat syntaxin-1A SNARE motif (residues 191–253), human fragments of SNAP-25A encoding its SNARE motifs (residues 11–82, referred to as SNAP-25_N, and residues 141–203, referred to as SNAP25_C; both SNAP-25 fragments including tryptophan at the N-terminus to facilitate detection by UV spectroscopy), rat complexin-1 fragment (residues 26–83), full-length rat complexin-1, and the MSP1E3D1 protein scaffold for nanodiscsc (18) were expressed and purified as described in (2). Full-length rat syntaxin-1A, full-length Cricetulus griseus NSF, and full-length Bos taurus αSNAP were expressed and purified as described in (19). The sequence of Syt1 C_2_AB fused through a 37-residue linker to the C-terminal SNARE motif of SNAP-25A (residues 141–204) (C_2_AB-linker-SNAP25_C), as described in (20), was subcloned from Addgene plasmid #70057 (p045) into the pGEX-KG vector. Synaptotagmin-1 C_2_B domain (residues 271–421) and C_2_AB fragment (residues 140–421) were expressed and purified as in (21, 22). As described in these studies, the following steps are necessary to achieve best purity of the Syt1 C_2_B domain and C_2_AB domain: treatment of the lysate with protamine sulfate, washes on the affinity column with CaCl_2_, and cation exchange chromatography in buffer containing CaCl_2_. Only the last peak of the cation exchange chromatogram contains the pure Syt1 fragments devoid of polyacidic contaminants. Following cation exchange, the last step of purification Syt1 C_2_B domain and C_2_AB fragment was size exclusion on Superdex 75 column (GE 16/60) in 20 mM HEPES pH 7.4 100 mM KCl 1 mM TCEP, unless specified otherwise. Syt1 C_2_AB-linker-SNAP25_C was purified analogously to the C_2_AB fragment but, because the flexible linker is prone to degradation, the purification had to be performed quickly and at cold temperatures to minimize degradation.

Expression vectors for mutant proteins were generated using standard PCR-based techniques with custom designed primers. These mutants included: Syt1 C_2_B (residues 271–421) E295K/Y338A, Y338D/A402T, R322E/K325E/R281Q, R322E/K325E/E295A, R322E/K325E/Y338D, R322E/K325E/Y338W, R322E/K325E/A402T; Syt1 C_2_AB (residues 140–421) T285W; C_2_AB-linker-SNAP25_C C277A/E412C, C277A/S391C; SNAP-25_N (residues 11–82) E27C, Q34C and R59C. All mutant proteins were purified as the WT proteins, including 0.5 mM TCEP in the final purification step for cysteine containing proteins. Isotopically labeled and perdeuterated Syt1 C_2_B domains were expressed using M9 expression media in 99.9% D2O with D-glucose (1,2,3,4,5,6,6-D7, 97–98%) as the sole carbon source (3 g/L) and ^15^NH_4_Cl as the sole nitrogen source (1 g/L). Specific ^13^CH_3_-labeling of the Met and Ile δ1 methyl groups of Syt1 C_2_B domains was achieved by adding [3,3–^2^H] ^13^C-methyl α-ketobutyric acid (80 mg/L) and ^13^C-methyl methionine (250 mg/L) (Cambridge Isotope Laboratories) to the cell cultures 30 min prior to IPTG induction. Syt1 C_2_B R322E/K325E/R281A was the only mutant used for NMR spectroscopy that was expressed without perdeuteration and methyl labelling. All C_2_B, C_2_AB and C_2_AB-linker-SNAP25_C fusion proteins were expressed overnight at 25 °C with 0.4 mM IPTG induction at OD 0.8.

#### NMR spectroscopy

All NMR spectra were acquired at 25 °C on Agilent DD2 spectrometers equipped with cold probes operating at 600 or 800 MHz. Titrations of WT and mutant ^2^H,^15^N-labeled Syt1 C_2_B domain specifically ^13^CH_3_-labeled at the Ile δ1 and Met methyl groups (referred to as ^15^N-C_2_B for simplicity) with SNARE complex four-helix bundle bound to a fragment spanning residues 26-83 of complexin-1 (referred to as CpxSC) were performed as described in (2). Specific ^13^CH_3_ labeling was performed for acquisition of ^1^H-^13^C heteronuclear multiple quantum (HMQC) spectra with these samples, although no such spectra are described here. Because the R322E/K325E mutation increases the stability of the C_2_B domain and slows down H/D exchange (2, 21), some amide groups in the β-strands are not fully protonated after expression in D_2_O and purification in buffers containing H_2_O, resulting in signal loss for the corresponding peaks. All newly purified C_2_B mutants bearing the R322E/K325E mutation were incubated at RT for one week or at 37 °C for 15 hours to facilitate full exchange, but some amide groups were still not fully exchanged after this procedure. The titrations were performed in NMR buffer containing 20 mM HEPES (pH 7.4), 100 mM KCl, 1 mM EDTA, 1 mM TCEP, 10% D_2_O and protease inhibitor cocktail [which contained 1 μM Antipain Dihydrochloride (Thermo Fischer Scientific: 50488492), 20 μM Leupeptin (Gold Bio: L01025) and 0.8 μM Aprotinin (Gold Bio: A655100)]. A ^1^H-^1^5N TROSY-HSQC spectrum was acquired first for 32 μM isolated ^15^N-C_2_B domain and additional ^1^H-^15^N TROSY-HSQC spectra were acquired after adding increasing concentrations of CpxSC to the sample, resulting in gradual dilution of the ^15^N-C_2_B domain. The protein concentrations of each titration step for each mutant are indicated in the legends of Fig. 4 and S15. Soluble SNARE complex was assembled as described in (2), concentrated at room temperature to a concentration above 250 μM using a 30 kDa centrifugation filter (Amicon) and exchanged into NMR buffer using Zeba Spin Desalting Columns, 7K MWCO, 10 mL (Thermo Fisher). Complexin-1 (26–83) was also concentrated above 250 μM using a 3 kDa centrifugation filter (Amicon) and exchanged into NMR buffer. SNARE-Complexin-1 (26–83) complex was preassembled with 20% excess Complexin-1 (26–83) before mixing with ^15^N-C2B domain. Total acquisition times ranged from 3.5 to 87.5 hr, depending on the sensitivity of the spectra. All NMR data were processed with NMRPipe (23) (Delaglio et al., 1995) and analyzed with NMRViewJ (24).

#### Labeling proteins with fluorescent tags

SNAP-25_N R59C mutant was labeled with Monobromobimane (mBBr), Thermo Fisher Scientific (M20381). mBBr delivered in powder was solubilized using DMSO to 10 mM stock concentration freshly before labeling reaction. After affinity tag cleavage, SNAP-25_N R59C was treated with 10X molar excess DTT and subjected to size exclusion chromatography (SEC) on a Superdex 75 column (GE 16/60) in labeling buffer (20 mM HEPES pH 7.4 125 mM KCl). Labelling was performed in a solution of 72 μM protein with 10X molar excess mBBr overnight at 4°C while rotating, and excess dye was removed by SEC.

SNAP-25_N Q34C and E27C mutants and C2AB-linker-SNAP25_C single cysteine mutants were labeled with the photostable Trolox-Cy3 and Trolox-Cy5 derivatives (25) conjugated with maleimide, respectively (LD550-MAL and LD650-MAL, Lumidyne Technologies). Lumidyne dyes delivered in powder were solubilized using DMSO to 10 mM stock concentration and stored in 10 μL aliquots at -80°C after solubilization. After cation exchange purification, C2AB-linker-SNAP25_C single cysteine mutants were quickly buffer exchanged to 20 mM HEPES pH 7.4 100 mM KCl 0.5 mM TCEP using a PD-10 column. For the SNAP-25_N single cysteine mutants the last purification step after affinity tag cleavage was size exclusion chromatography on a Superdex 75 column (GE 16/60) with the labeling buffer 20 mM HEPES pH 7.4 100 mM KCl 0.5 mM TCEP. The buffer exchanged proteins at a concentration of 50–100 μM were incubated with 3X molar excess dye overnight at 4°C. The dye was added gently, preferably while stirring to avoid high local concentrations of DMSO. The reaction was quenched by addition of 10 mM DTT. Unreacted dye was separated from labeled protein by passing through a PD-10 column followed by size exclusion chromatography on a Superdex 75 column (GE 16/60). The concentration and the labeling efficiency of the tagged proteins were determined using UV-vis spectroscopy.

#### Bimane fluorescence quenching assay

All fluorescence emission scans were collected on a PTI Quantamaster 400 spectrofluorometer (T-format) at room temperature with slits set to 1.25 mm. For tryptophan-induced bimane fluorescence quenching assays, we used SNARE complexes anchored on nanodiscs as described (2) and labeled with bimane at position R59C of SNAP-25_N. The lipid composition of nanodiscs was 84% POPC, 15% DOPS, 1% PIP_2_. The experiments were performed in 25 mM HEPES pH 7.4 100 mM KCl 0.1 mM TCEP 2.5 mM MgCl_2_, 2 mM ATP 1 mM EGTA containing 1.5 μM BSA to prevent sample binding to the cuvette. Fluorescence emission spectra (excitation at 380 nm) were acquired for samples containing 1 μM SC-59-bimane-nanodiscs alone or in the presence of 4 μM C_2_A-T285W without or with 2.0 mM CaCl_2_ (1.0 mM free Ca^2+^).

#### Preparation of C_2_AB-SNARE-Complex-liposomes

To prepare liposomes containing C_2_AB covalently linked to the SNARE complex, we used a lipid mixture including 84% POPC, 15% DOPS, 1% PIP2. Lipids were blow-dried under a stream of nitrogen, tilting and rotating the tube to form a thin film on the wall of the tubes. If precipitation was observed in the lipid mixture due to PIP_2_ addition, the tube was incubated in 45 °C until solution cleared before drying. The tube with lipid film was then incubated in the vacuum desiccator overnight for further drying. Detergent solubilized C_2_AB-SNARE complex was formed by incubating 0.25 μM Trolox-Cy3-labeled SNAP-25_N E27C or Q34C with 0.3 μM full-length rat syntaxin-1A, 0.3 μM synaptobrevin-2 (29–93) and 0.3 μM Trolox-Cy5-labeled C_2_AB-linker-SNAP25_C S391C or E412C in the presence of 1% β−OG, 0.5 M NaCl and protease inhibitor cocktail overnight at 4°C. Next day, 25mM HEPES pH 7.4 100mM KCl buffer containing 2% (wt/vol) β-OG was added to hydrate the lipid film. The tube was vortexed vigorously for complete suspension (∼2 min). The lipids were incubated at room temperature for 15 min and afterward sonicated in a bath sonicator for 5 min twice. After confirming formation of SDS-resistant C2AB-SNARE-complex by the SDS-PAGE, the complex was mixed with solubilized lipids in protein to lipid (P:L) ratio 1:5000 in the presence of 1% β−OG and incubated for 25 min at RT. The sample was dialyzed using the Thermo Scientific™ Slide-A-Lyzer™ Dialysis Cassettes MWCO 10kDa against 25mM HEPES pH 7.4 100mM KCl buffer containing Amberlite XAD-2 detergent-absorbing beads. The dialysis was started in 0.5 L buffer with 1 g XAD-2 and incubated while mixing with a magnetic stirrer for 1 h in 4°C, changed to 0.5 L fresh buffer with 1 g XAD-2 and incubated for 1 h, and then changed again to 1 L fresh buffer with 2 g XAD-2 and dialyzed over-night (∼12 h). The proteoliposomes were always kept on ice and used for the fluorescence experiments. Size distribution analysis by dynamic light scattering showed the average liposome radii was typically ∼60–95 nm.

#### FRET assays

All fluorescence emission scans were collected on a PTI Quantamaster 400 spectrofluorometer (T-format) at room temperature with excitation at 550 nm and slits set to 1.25 mm. Each sample contained 0.125 μM C2AB-SNARE-complex-liposomes in 25 mM HEPES pH 7.4 100 mM KCl buffer, 1 mM EGTA, 2.5 mM MgCl2, 2 mM ATP, 1.5 μM BSA, and 0.15 μM full-length Complexin-1. Mg-ATP and complexin-1 were included to hinder non-specific interactions. To examine the reproducibility of Ca^2+^-induced changes in FRET, spectra were acquired for separate samples prepared under identical conditions before and after addition of 2.0 mM CaCl_2_ (1.0 mM free Ca^2+^). Because the Syt1 C_2_AB fragment and SNARE complex were fused covalently via the 37 aa linker and the SNARE complex is resistant to SDS, control spectra to measure the maximum donor fluorescence observable without FRET were acquired after incubation for 5 minutes at 37 oC with 0.4 µM NSF and 2 µM αSNAP (to disassemble the SNARE complex) in 25 mM HEPES pH 7.4 100 mM KCl buffer containing 1 mM EGTA, 2.5 mM MgCl_2_, 2 mM ATP, 1.5 μM BSA and 1% BOG. The presence of detergent proved to be necessary to recover maximum signal because a subset of the C_2_AB-SNARE-complexes get reconstituted inside the liposomes and hence are inaccessible to the disassembly machinery.

---

**Table S1.** Parameters of the MD simulations^a^.

| *Vesicle* | CHL1 | POPC | SAGL | SAPE | SAPI2D | SDPE | SDPS | SOPS | total |
| --- | --- | --- | --- | --- | --- | --- | --- | --- | --- |
| Outer leaflet | 1258 | 296 | 0 | 534 | 0 | 282 | 210 | 199 | 2779 |
| % | 45.3 | 10.6 | 0 | 19.2 | 0 | 10.1 | 7.6 | 7.2 |  |
| Inner leaflet | 814 | 668 | 0 | 183 | 0 | 99 | 1 | 1 | 1766 |
| % | 46.1 | 37.8 | 0 | 10.4 | 0 | 5.6 | 0.1 | 0.1 |  |
| *Flat bilayer 1* | CHL1 | POPC | SAGL | SAPE | SAPI2D | SDPE | SDPS | SOPS | total |
| Upper leaflet | 830 | 151 | 18 | 132 | 93 | 262 | 182 | 181 | 1849 |
| % | 44.9 | 8.2 | 1.0 | 7.1 | 5.0 | 14.2 | 9.8 | 9.8 |  |
| Lower leaflet | 810 | 774 | 18 | 72 | 0 | 126 | 0 | 0 | 1800 |
| % | 45.0 | 43.0 | 1.0 | 4.0 | 0 | 7.0 | 0 | 0 |  |
| *Flat bilayer 2* | CHL1 | POPC | SAGL | SAPE | SAPI2D | SDPE | SDPS | SOPS | total |
| Upper leaflet | 748 | 130 | 17 | 115 | 82 | 231 | 165 | 161 | 1649 |
| % | 45.3 | 7.9 | 1.0 | 7.0 | 5.0 | 14.0 | 10.0 | 9.8 | 100 |
| Lower leaflet | 720 | 688 | 16 | 64 | 0 | 112 | 0 | 0 | 1600 |
| % | 45.0 | 43.0 | 1.0 | 4.0 | 0 | 7.0 | 0 | 0 | 100 |
| *Flat bilayer 3* | CHL1 | POPC | SAGL | SAPE | SAPI2D | SDPE | SDPS | SOPS | total |
| Upper leaflet | 1035 | 184 | 23 | 161 | 115 | 322 | 230 | 230 | 2300 |
| % | 45 | 8 | 1 | 7 | 5 | 14 | 10 | 10 |  |
| Lower leaflet | 1009 | 964 | 22 | 88 | 0 | 154 | 0 | 0 | 2237 |
| % | 45.1 | 43.1 | 1.0 | 3.9 | 0.0 | 6.9 | 0.0 | 0.0 |  |
| system name | flat bilayer | number of atoms | initial cell dimensions |  |  | length and temperature |  |  |  |
| cac2absc | 1 | 4921862 | 36.9 x 36.4 x 36.7 nm<sup>3</sup> |  |  | 668 ns 310 K |  |  |  |
| fusiong | 1 | 4922564 | 36.9 x 36.4 x 36.7 nm<sup>3</sup> |  |  | 510 ns 310 K |  |  |  |
| nosytfusion | 1 | 4921880 | 36.9 x 36.4 x 36.7 nm<sup>3</sup> |  |  | 510 ns 310 K |  |  |  |
| sytfusion2g | 1 | 4920453 | 36.9 x 36.4 x 36.7 nm<sup>3</sup> |  |  | 570 ns 310 K |  |  |  |
| sytfusion3 | 2 | 4586690 | 35.6 x 36.8 x 35.3 nm<sup>3</sup> |  |  | 450 ns 310 K |  |  |  |
| s1action2 | 3 | 6745506 | 42.5 x 37.5 x 42.5 nm<sup>3</sup> |  |  | 596 ns 310 K |  |  |  |

<sup>a</sup>All systems included four trans-SNARE complexes, a 24 nm vesicle and the indicated flat bilayer. The cac2absc, fusiong, sytfusion3 and s1action2 had four C<sub>2</sub>AB molecules, sytfusion2g had two and nosytfusion had none. C<sub>2</sub>AB molecules had five bound Ca<sup>2+</sup> ions except for those in s1action2, which was Ca<sup>2+</sup> free.

**Table S2.**
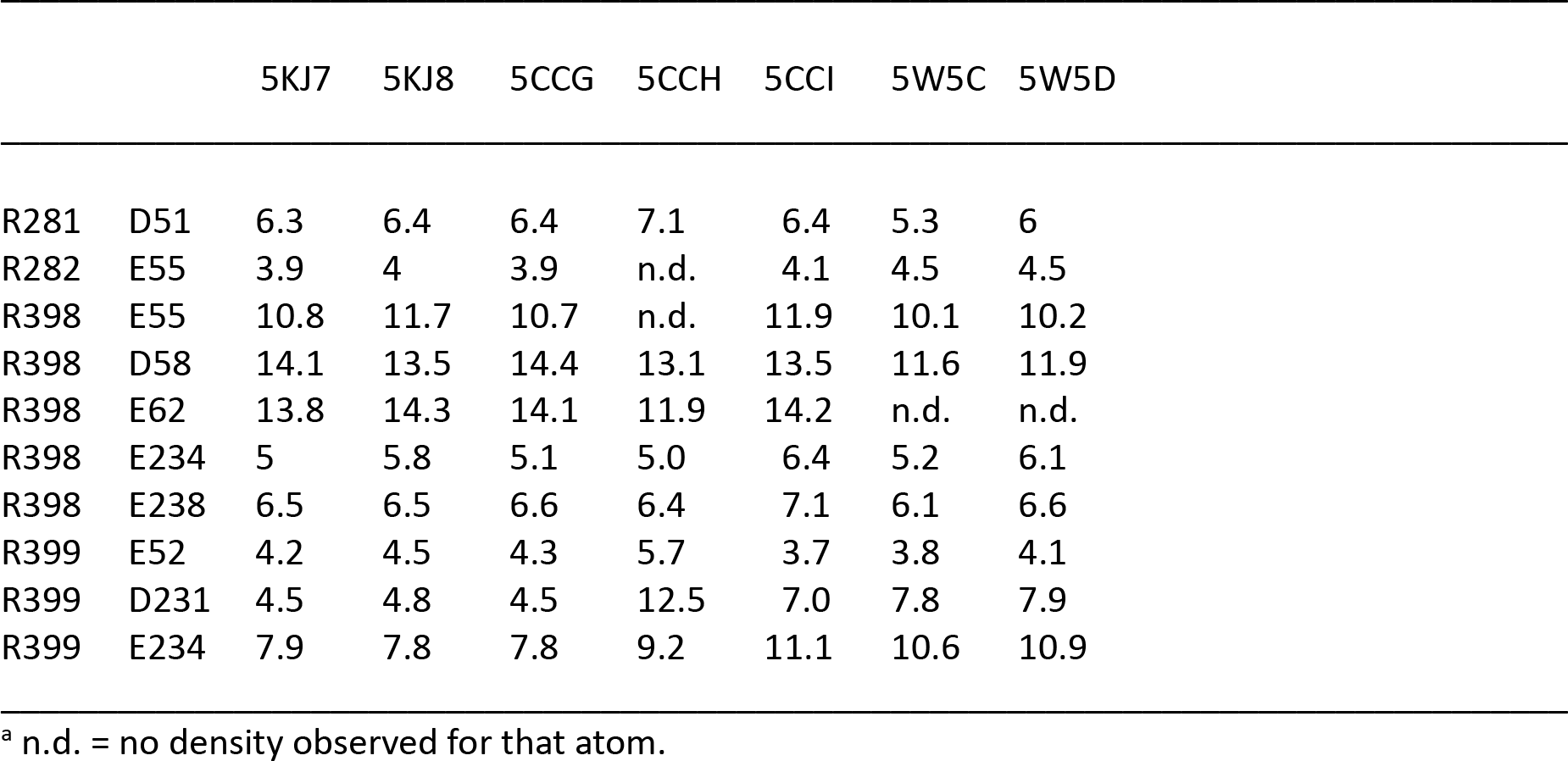
Distances (Å) between the CZ atoms of arginines in region II of the primary interface and CG atoms of aspartes or CD atoms of glutamates of the SNAREs in crystal structures of Syt1-SNARE complexes^a^.

**Table S3.** Average distances (Å) between the CZ atoms of arginines in region II of the primary interface and CG atoms of aspartes or CD atoms of glutamates of the SNAREs in the four primed complexes (prcs) during the s1action2 MD simulation, and average distances for the four complexes of s1action 2 and the four complexes of the prsg MD simulation of ref. (10).

|  |  | prc-1 | prc-2 | prc-3 | prc-4 | S1action2<br>average | prsg<br>average |
| --- | --- | --- | --- | --- | --- | --- | --- |
| R281 | D51 | 4.1 | 4.0 | 4.1 | 3.9 | 4.0 | 4.0 |
| R282 | E55 | 4.5 | 4.6 | 4.3 | 4.5 | 4.5 | 4.5 |
| R398 | E55 | 6.8 | 6.7 | 7.9 | 6.8 | 7.0 | 5.7 |
| R398 | D58 | 7.6 | 7.6 | 10.0 | 7.2 | 8.1 | 6.0 |
| R398 | E62 | 5.7 | 4.4 | 8.3 | 4.9 | 5.8 | 5.9 |
| R398 | E234 | 8.3 | 9.4 | 7.2 | 9.1 | 8.5 | 10.9 |
| R398 | E238 | 9.0 | 10.4 | 7.3 | 10.4 | 9.3 | 17.0 |
| R399 | E52 | 5.7 | 8.4 | 5.2 | 5.6 | 6.2 | 5.5 |
| R399 | D231 | 7.0 | 9.1 | 7.3 | 7.2 | 7.6 | 6.6 |
| R399 | E234 | 6.3 | 6.3 | 8.7 | 7.1 | 7.1 | 8.1 |

**Figure S1.**
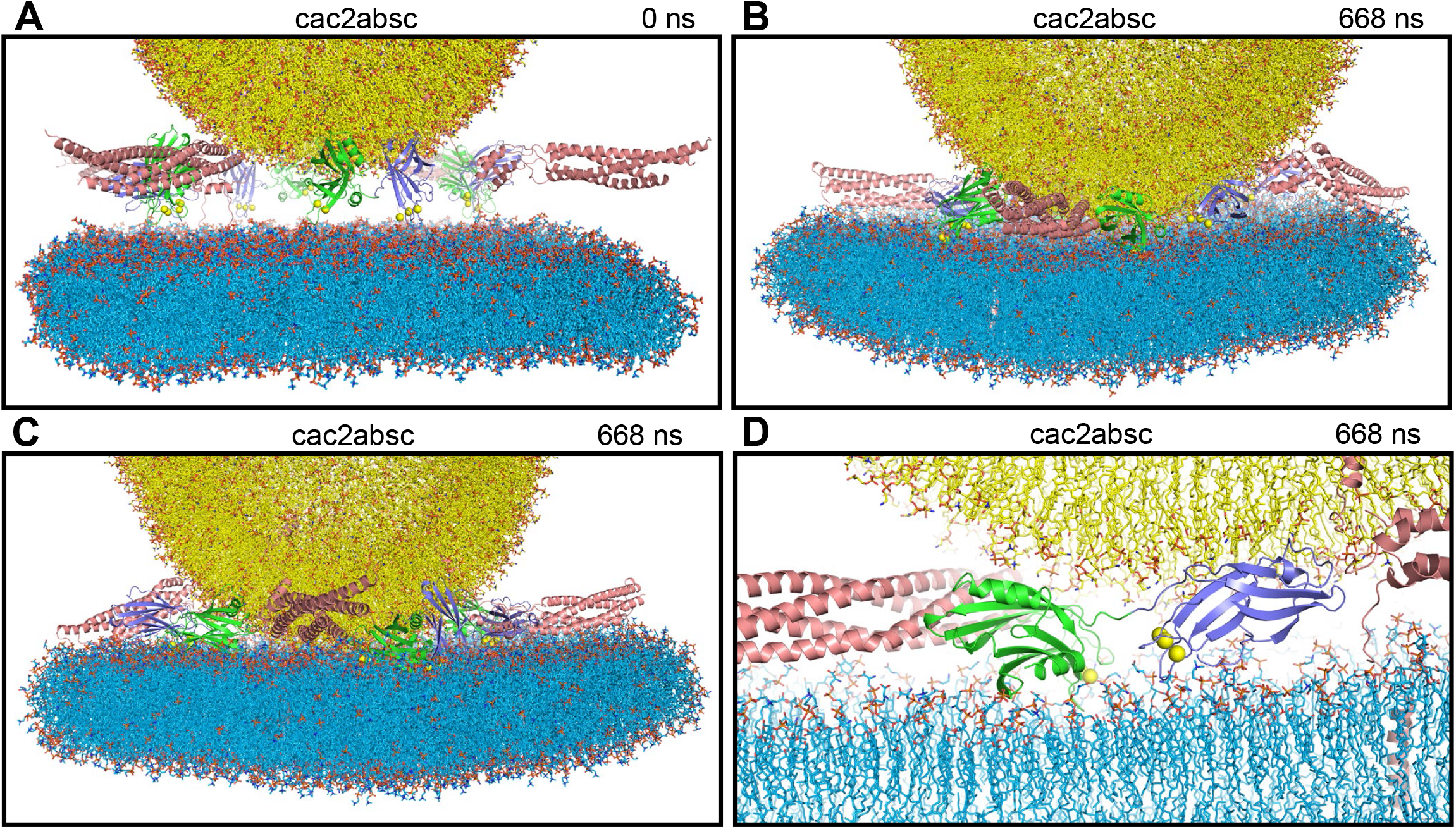
MD simulation designed to investigate whether Ca^2+^-saturated Syt1 C_2_ domains spontaneously insert into the bilayers, perturb or bridge the bilayers (cac2absc simulation). (*A*) Initial configuration. (*B, C*) Configuration after 668 ns of simulation viewed from opposed angles. (*D*) Close-up view of the system at 668 ns showing a C_2_B domain that is next to the C-terminus of a SNARE complex that is almost fully assembled and with the Ca^2+^-binding loops oriented toward the center of the membrane-membrane interface. Lipids are represented by stick models, proteins by ribbon diagrams and Ca^2+^ ions by spheres with the same color coding as in Fig. 1.

**Figure S2.**
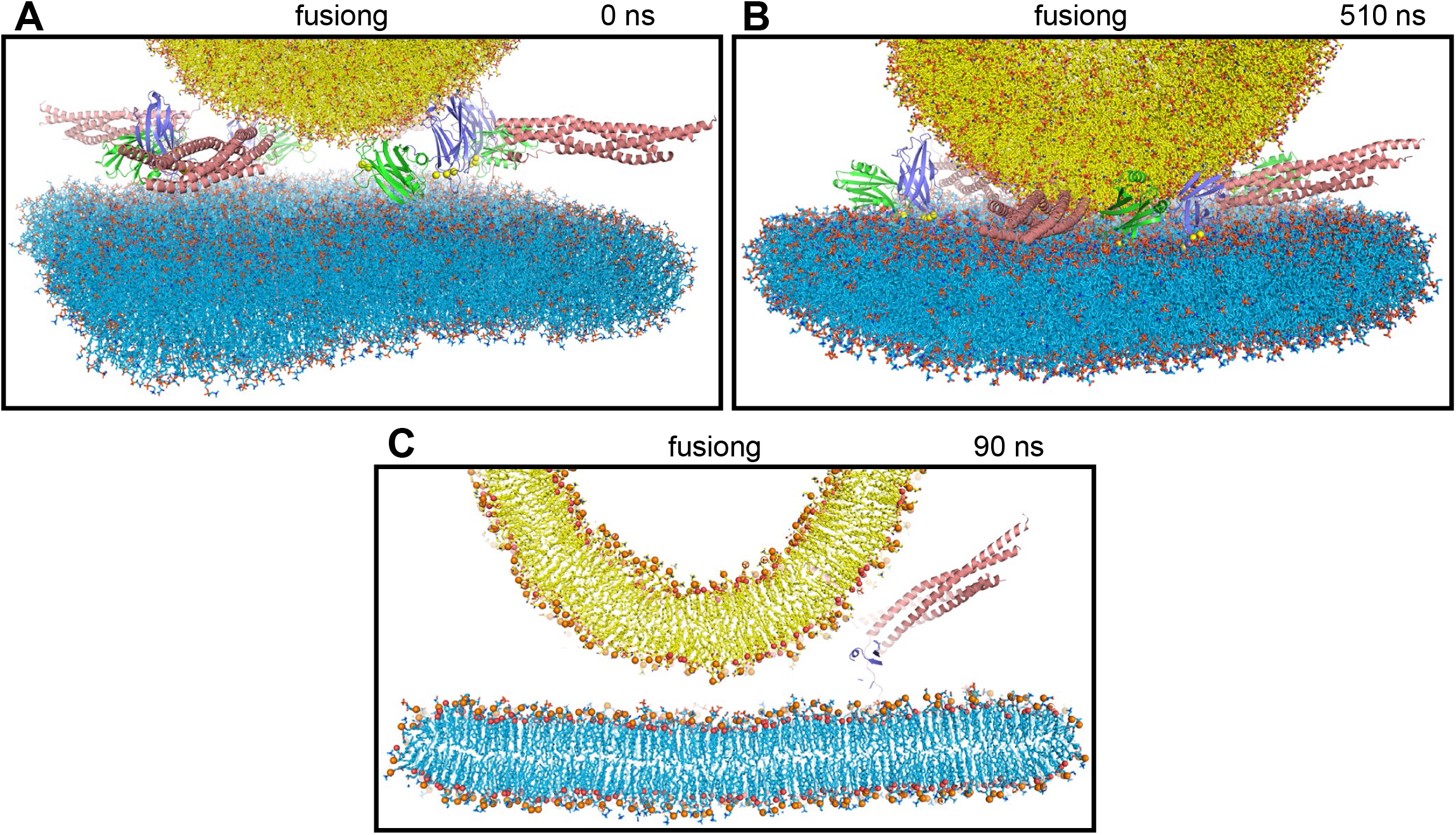
MD simulation designed to examine whether the Ca^2+^-saturated Syt1 C_2_ domains cooperate with almost fully assembled SNARE complexes to induce membrane fusion (fusiong simulation). (*A*) Initial configuration. (*B*) Configuration after 510 ns of simulation. (*C*) Slice of the system after 90 ns of simulation showing how the membrane remained more distant than in an analogous simulation without Syt1 C_2_AB molecules (compared with Fig. S3C). Lipids are represented by stick models, proteins by ribbon diagrams and Ca^2+^ ions by spheres with the same color coding as in Fig. 1.

**Figure S3.**
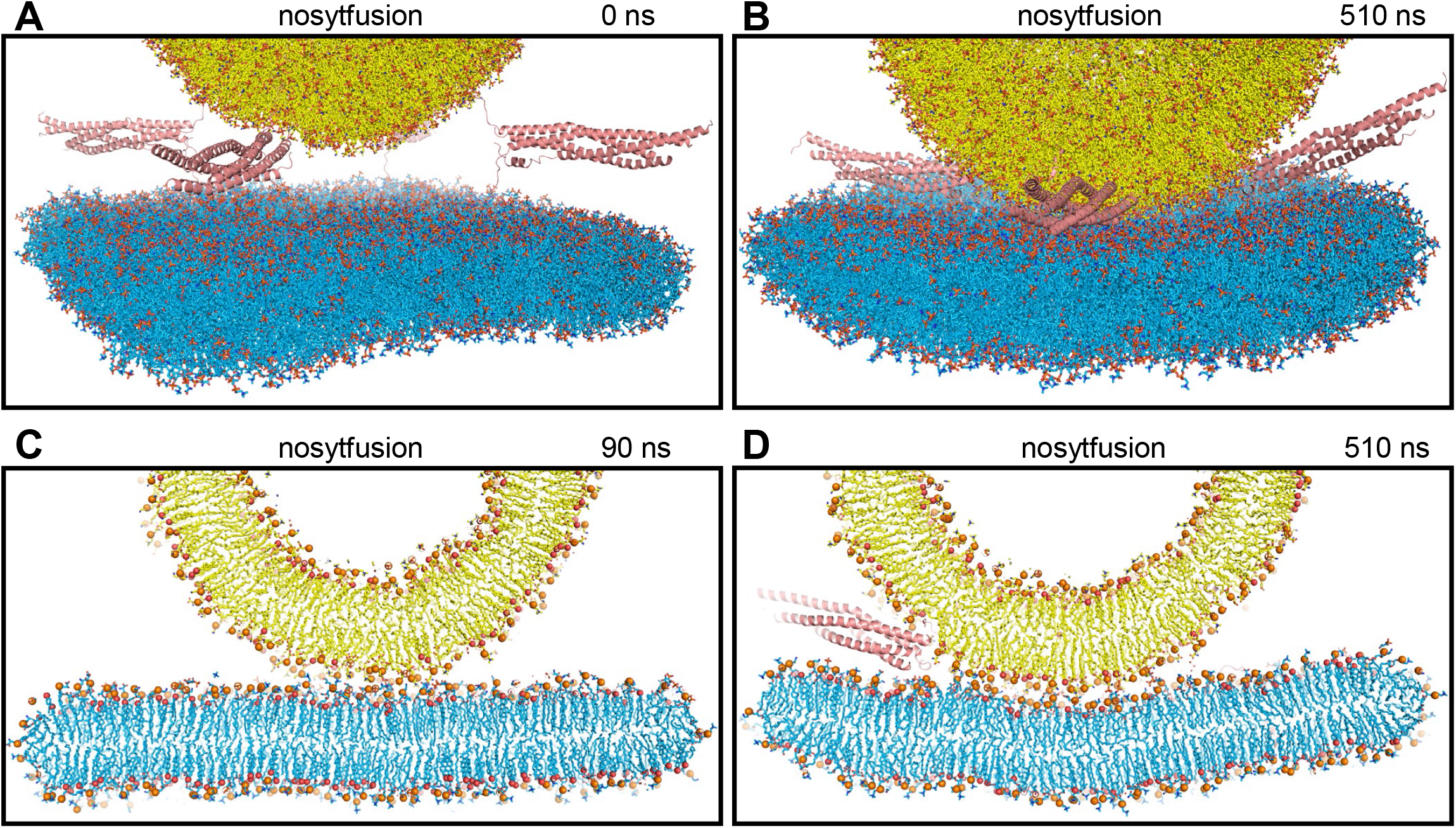
MD simulation of a system analogous to fusiong but without C_2_AB molecules to test whether they hinder the action of SNARE complexes in bringing membranes together (nosytfusion simulation). (*A*) Initial configuration. (*B*) Configuration after 510 ns of simulation. (*C, D*) Slices of the system after 90 ns (*C*) and 510 ns (*D*). Lipids are represented by stick models, proteins by ribbon diagrams and Ca^2+^ ions by spheres with the same color coding as in Fig. 1. Panel (*C*) shows how the membranes were already in contact at 90 ns, in contrast with the simulation that included C_2_AB molecules (Fig. S2C). Panel (*D*) shows that the two membranes started to form an extended interface, unlike the simulation including C_2_AB molecules (Fig. 1 b).

**Figure S4.**
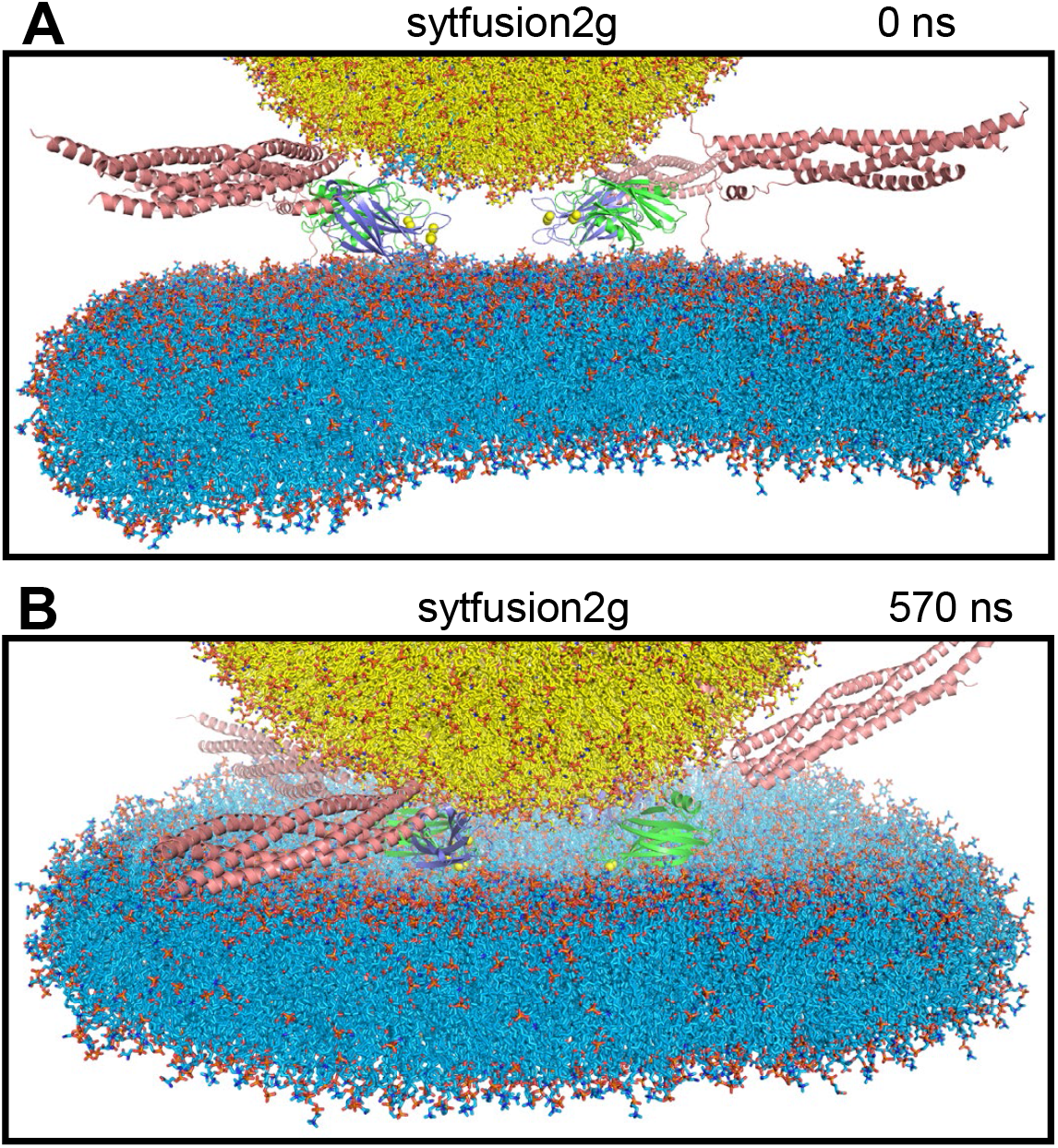
MD simulation with two Ca^2+^-saturated Syt1 C_2_ domains located between the two membranes to study whether they might play a direct role in membrane fusion (sytfusion2g simulation). Lipids are represented by stick models, proteins by ribbon diagrams and Ca^2+^ ions by spheres with the same color coding as in Fig. 1. (*A*) Initial configuration. (*B*) Configuration after 570 ns of simulation.

**Figure S5.**
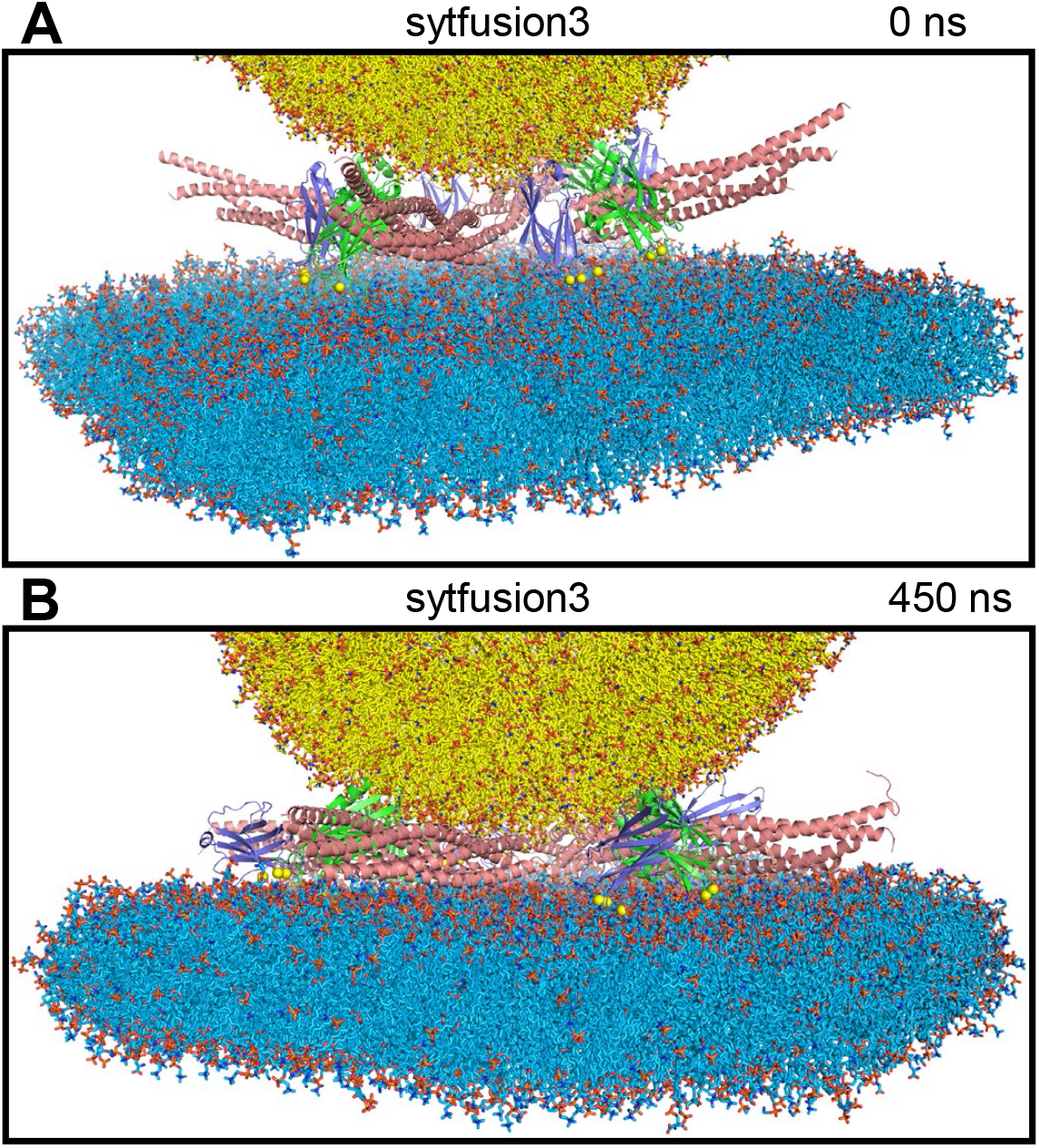
MD simulation designed to examine whether Ca^2+^-saturated Syt1 C_2_ domains might act as wedges that prevent the membranes from coming closer while the SNARE complexes pull the membranes together in the center to induce to torque forces that help to bend the membranes to initiate membrane fusion (sytfusion3 simulation). Lipids are represented by stick models, proteins by ribbon diagrams and Ca^2+^ ions by spheres with the same color coding as in Fig. 1. (*A*) Initial configuration. (*B*) Configuration after 450 ns of simulation.

**Figure S6.**
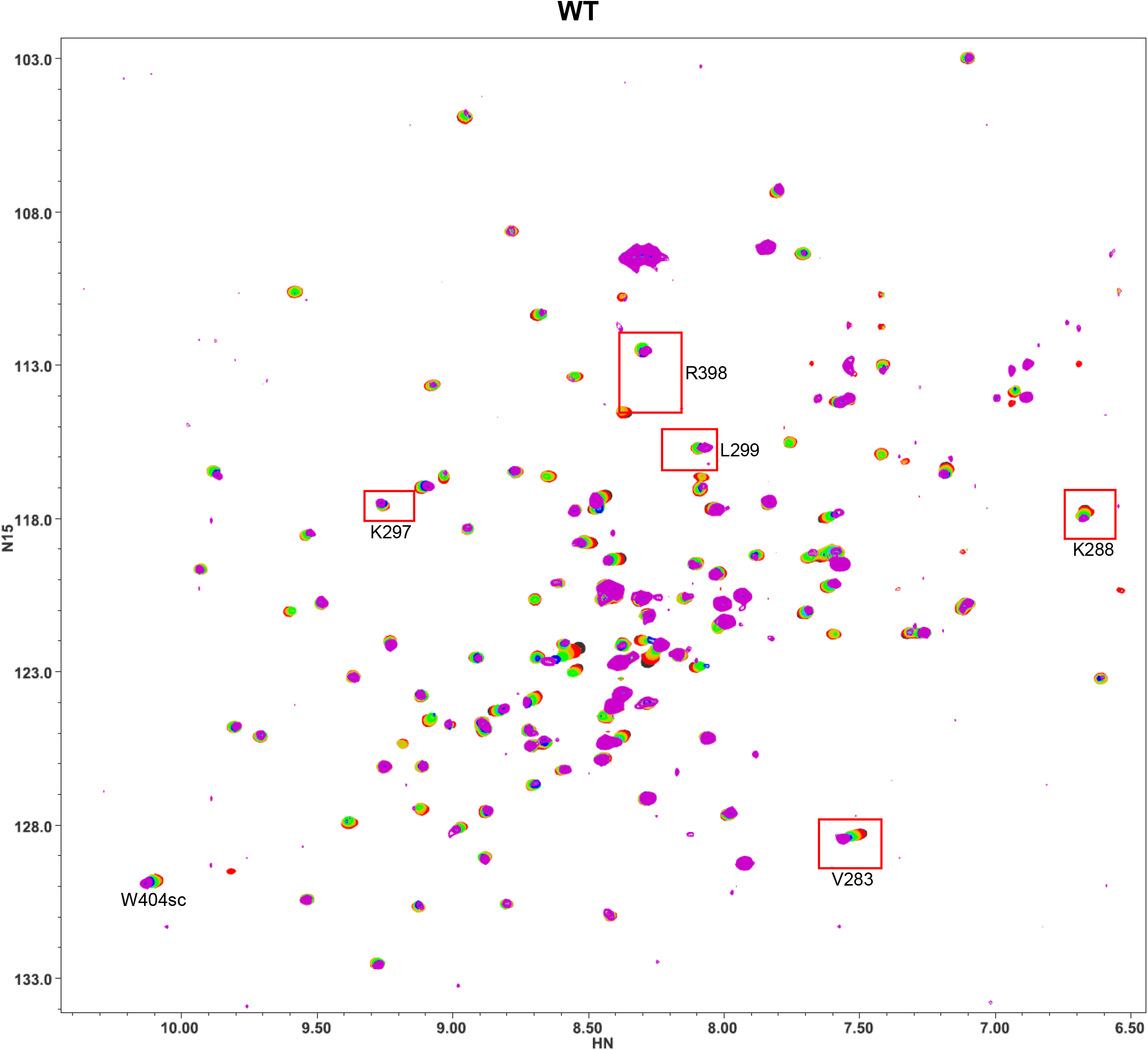
Superposition of ^1^H-^15^N TROSY-HSQC spectra of WT ^15^N-C_2_B domain in the absence of Ca^2+^ and the presence of increasing concentrations of CpxSC. The following concentrations of ^15^N-C_2_B and CpxSC (μM/μM) were used (from black to purple): 32/0, 30/10, 28/19, 26/28, 24/36, 20/51, 17/64, 12/85. The intensities of cross-peaks decreased as CpxSC was added because ^15^N-C_2_B was diluted and binding to CpxSC causes cross-peak broadening. Contour levels were adjusted to compensate for these decreased intensities, but some cross-peaks may not be visible even after these adjustments at the higher CpxSc concentrations. The red boxes indicate the cross-peaks that are displayed in the expansions of Fig. S15B.

**Figure S7.**
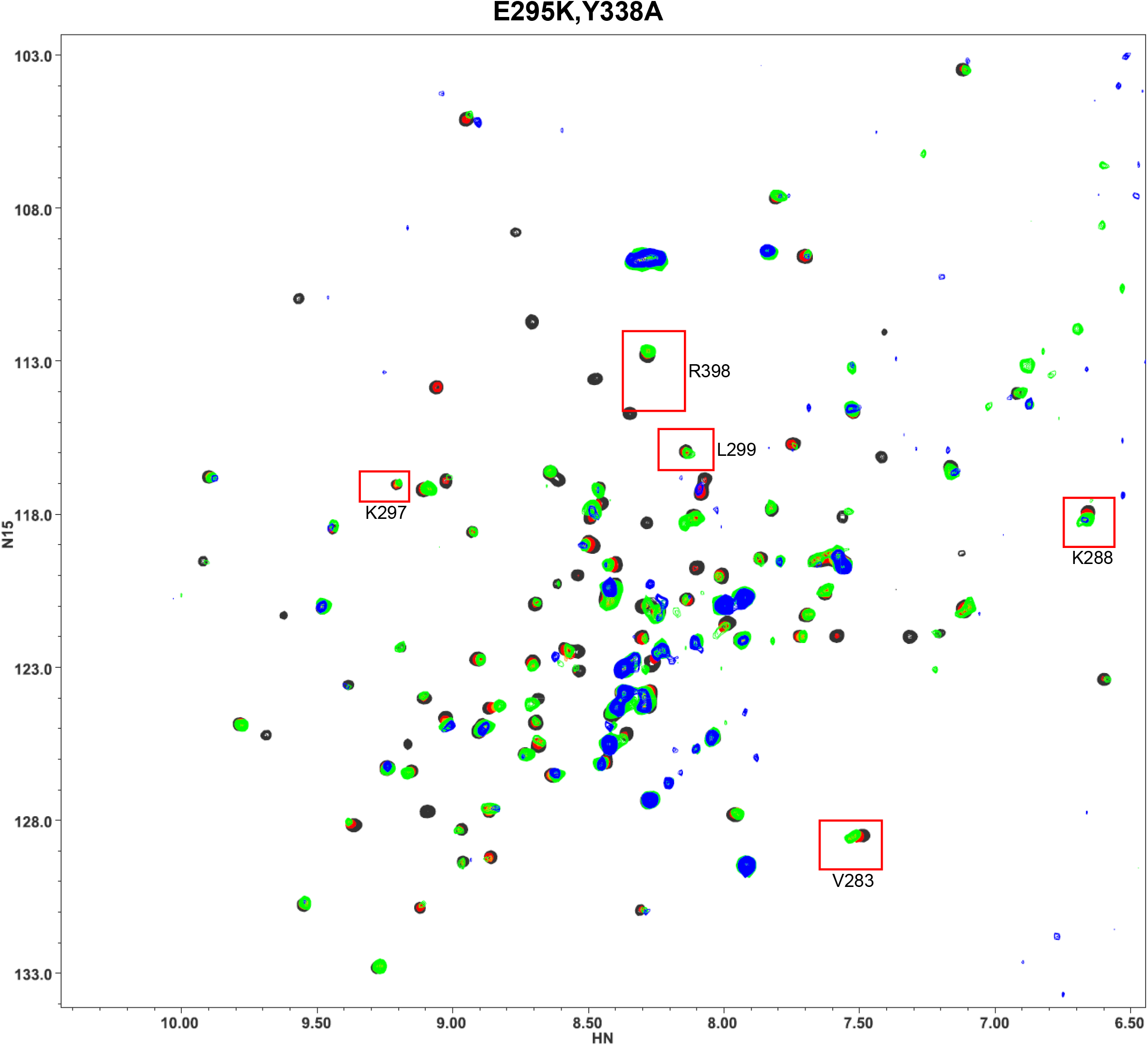
Superposition of ^1^H-^15^N TROSY-HSQC spectra of ^15^N-C_2_B domain E295K/Y338A mutant in the absence of Ca^2+^ and the presence of increasing concentrations of CpxSC. The following concentrations of ^15^N-C_2_B and CpxSC (μM/μM) were used (from black to blue): 32/0, 30/11, 27/21, 25/30, 23/38, 19/59. The intensities of cross-peaks decreased as CpxSC was added because ^15^N-C_2_B was diluted and binding to CpxSC causes cross-peak broadening. Contour levels were adjusted to compensate for these decreased intensities, but some cross-peaks may not be visible even after these adjustments at the higher CpxSc concentrations. The red boxes indicate the cross-peaks that are displayed in the expansions of Fig. S15B.

**Figure S8.**
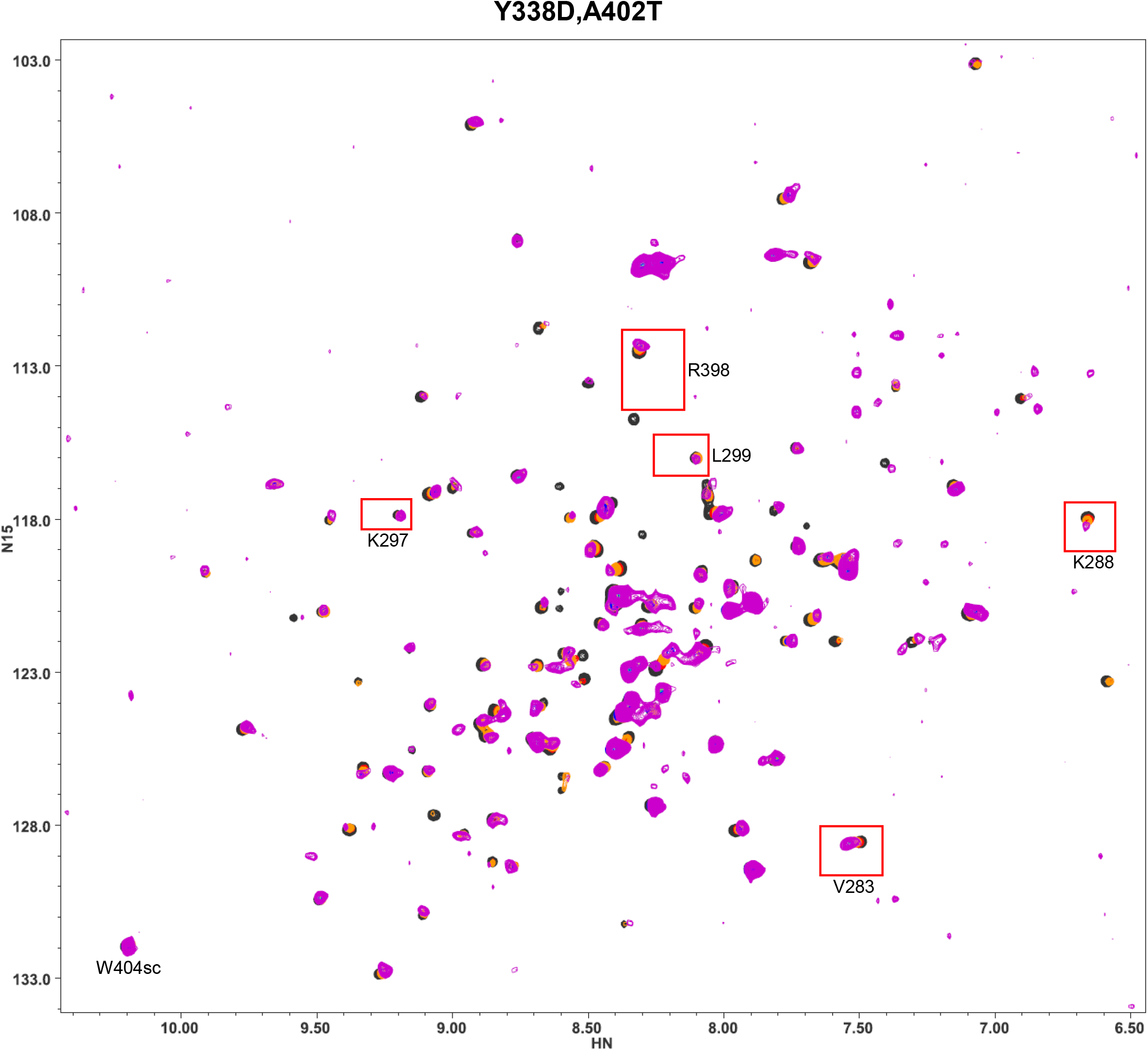
Superposition of ^1^H-^15^N TROSY-HSQC spectra of ^15^N-C_2_B domain Y338D/A402T mutant in the absence of Ca^2+^ and the presence of increasing concentrations of CpxSC. The following concentrations of ^15^N-C_2_B and CpxSC (μM/μM) were used (from black to purple): 32/0, 30/10, 28/19, 26/28, 24/36, 20/51, 17/64, 12/86. The intensities of cross-peaks decreased as CpxSC was added because ^15^N-C_2_B was diluted and binding to CpxSC causes cross-peak broadening. Contour levels were adjusted to compensate for these decreased intensities, but some cross-peaks may not be visible even after these adjustments at the higher CpxSc concentrations. The red boxes indicate the cross-peaks that are displayed in the expansions of Fig. S15B.

**Figure S9.**
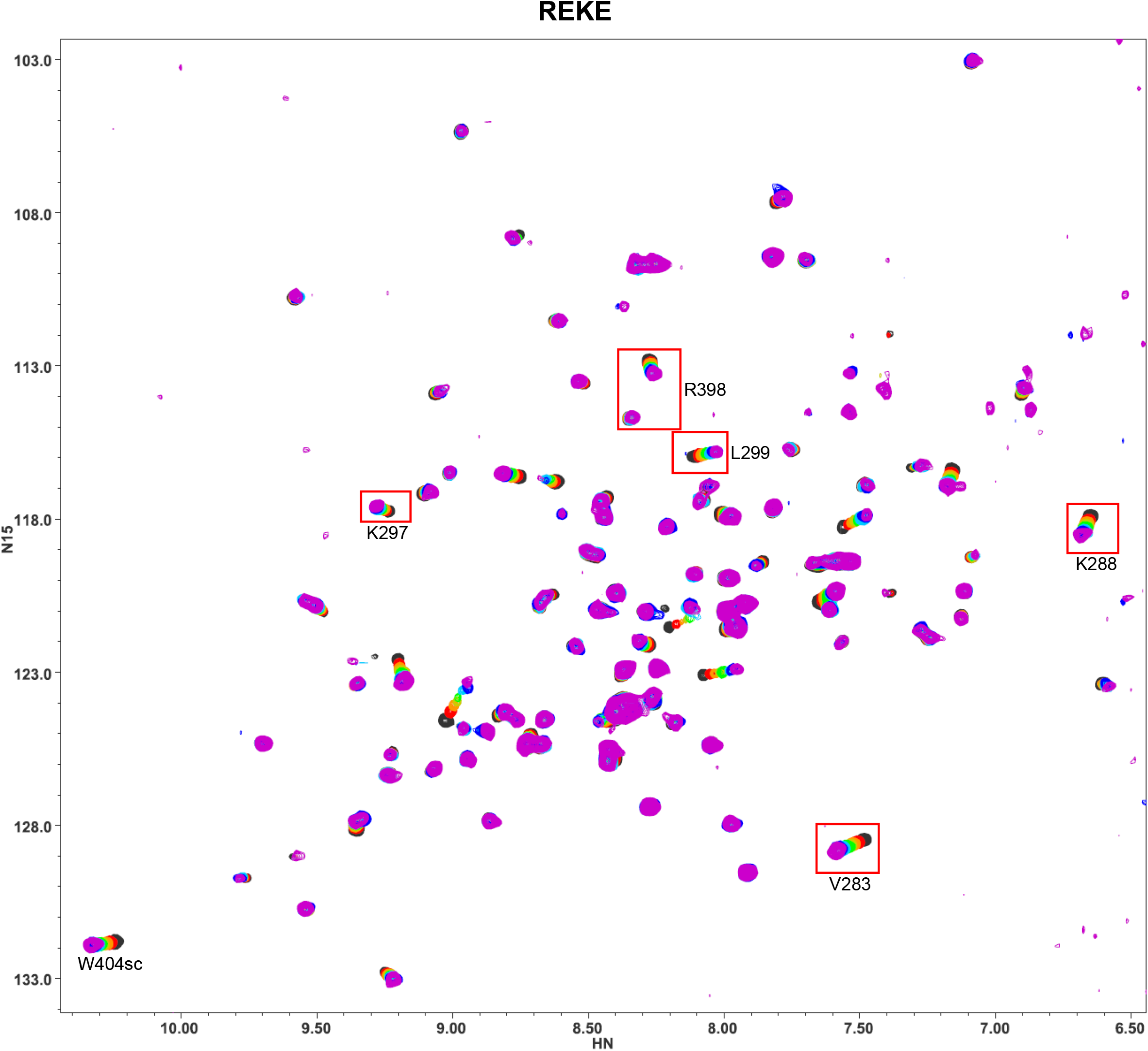
Superposition of ^1^H-^15^N TROSY-HSQC spectra of ^15^N-C_2_B domain REKE mutant in the absence of Ca^2+^ and the presence of increasing concentrations of CpxSC. The following concentrations of ^15^N-C_2_B and CpxSC (μM/μM) were used (from black to purple): 32/0, 30/10, 28/19, 26/28, 24/36, 20/51, 17/64, 12/85. The intensities of cross-peaks decreased as CpxSC was added because ^15^N-C_2_B was diluted and binding to CpxSC causes cross-peak broadening. Contour levels were adjusted to compensate for these decreased intensities, but some cross-peaks may not be visible even after these adjustments at the higher CpxSc concentrations. The red boxes indicate the cross-peaks that are displayed in the expansions of Fig. 4.

**Figure S10.**
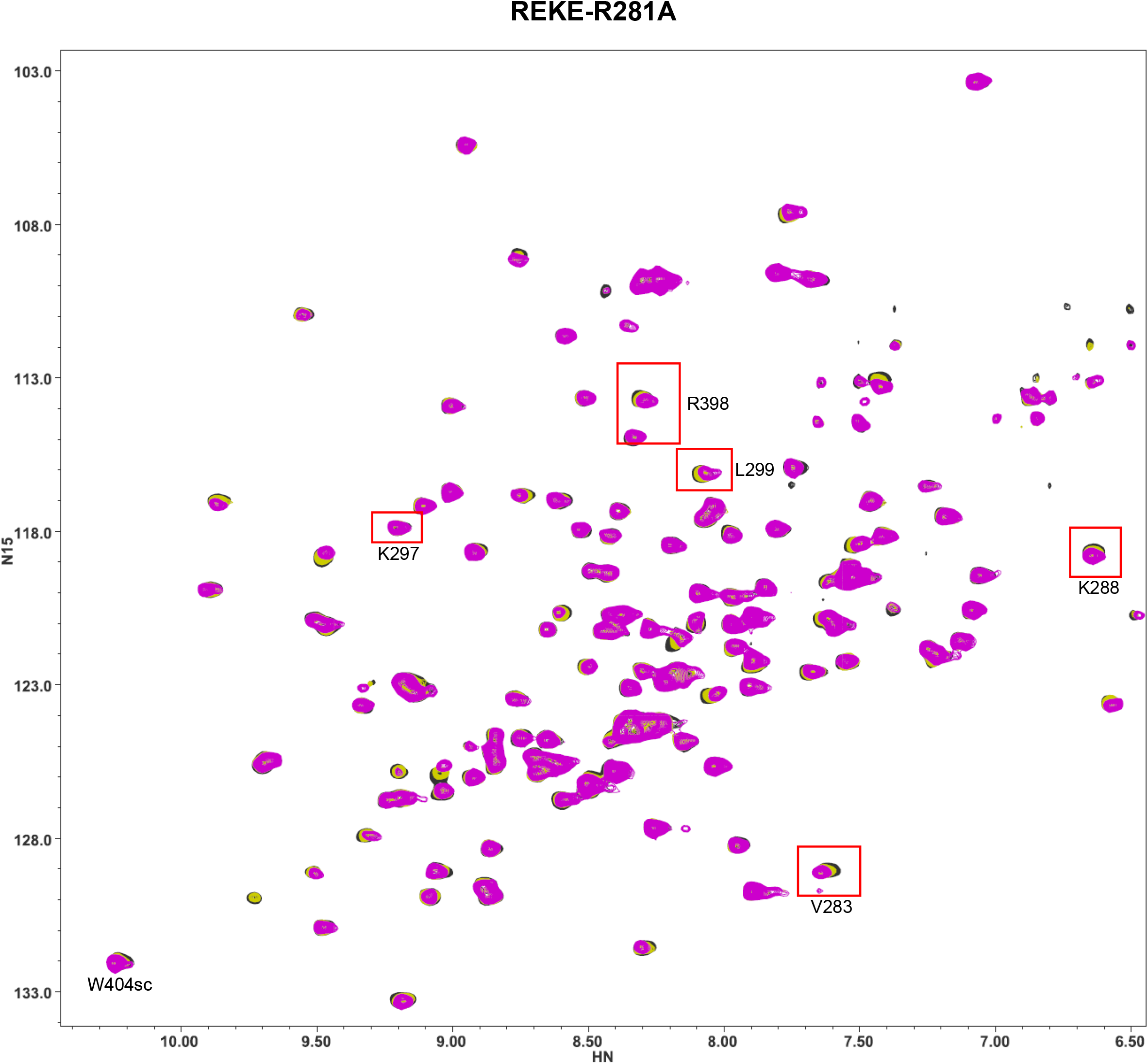
Superposition of ^1^H-^15^N TROSY-HSQC spectra of ^15^N-C_2_B domain REKE-R281A mutant in the absence of Ca^2+^ and the presence of increasing concentrations of CpxSC. The following concentrations of ^15^N-C_2_B and CpxSC (μM/μM) were used: 32/0 (black), 26/28 (yellow) and 13/98 (purple). The intensities of cross-peaks decreased as CpxSC was added because ^15^N-C_2_B was diluted and binding to CpxSC causes cross-peak broadening. Contour levels were adjusted to compensate for these decreased intensities, but some cross-peaks may not be visible even after these adjustments at the higher CpxSc concentrations. The red boxes indicate the cross-peaks that are displayed in the expansions of Fig. 4.

**Figure S11.**
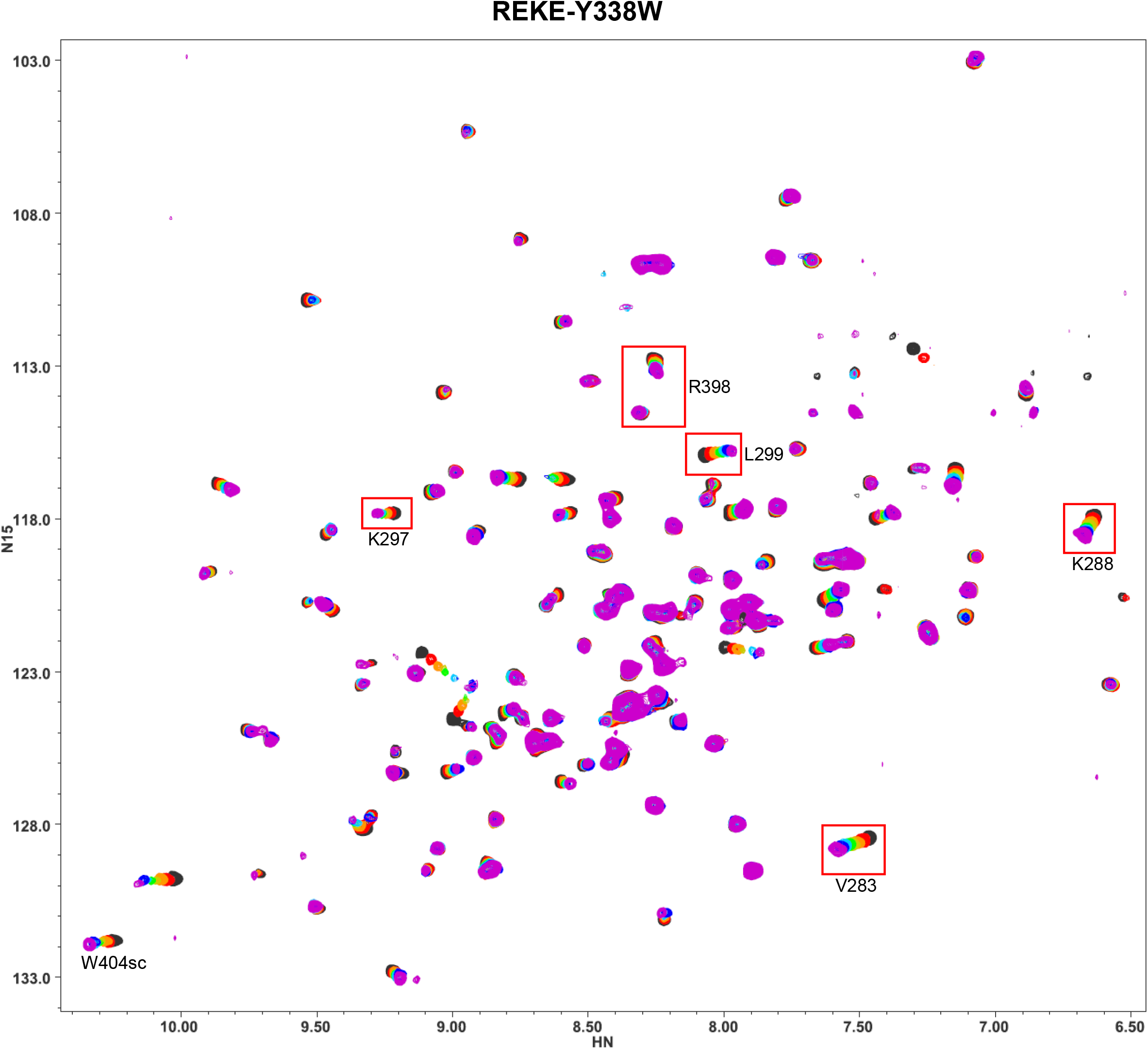
Superposition of ^1^H-^15^N TROSY-HSQC spectra of ^15^N-C_2_B domain REKE-Y338W mutant in the absence of Ca^2+^ and the presence of increasing concentrations of CpxSC. The following concentrations of ^15^N-C_2_B and CpxSC (μM/μM) were used (from black to purple): 32/0, 30/10, 28/19, 26/28, 24/36, 20/53, 17/67, 12/88. The intensities of cross-peaks decreased as CpxSC was added because ^15^N-C_2_B was diluted and binding to CpxSC causes cross-peak broadening. Contour levels were adjusted to compensate for these decreased intensities, but some cross-peaks may not be visible even after these adjustments at the higher CpxSc concentrations. The red boxes indicate the cross-peaks that are displayed in the expansions of Fig. 4.

**Figure S12.**
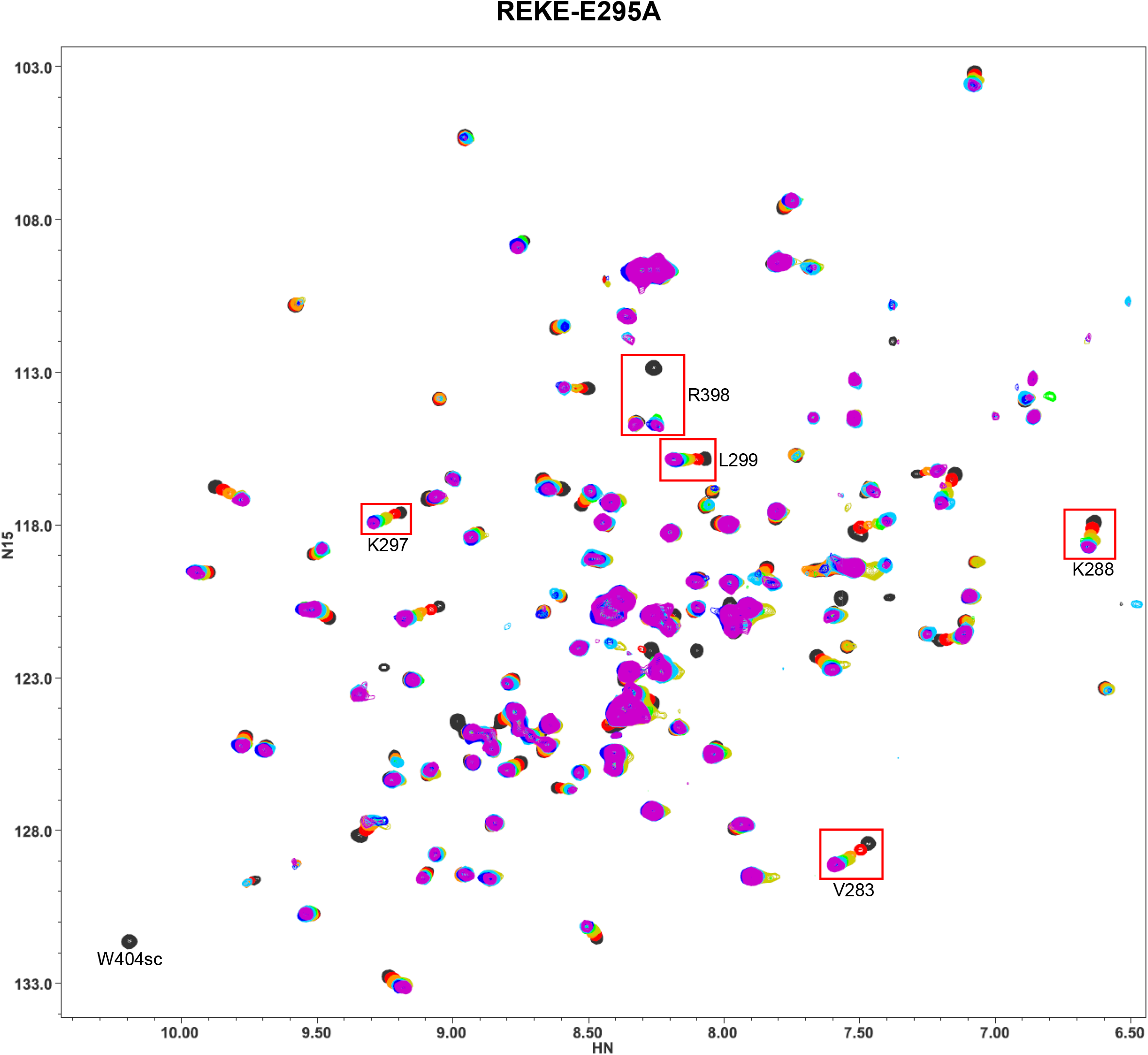
Superposition of ^1^H-^15^N TROSY-HSQC spectra of ^15^N-C_2_B domain REKE-E295A mutant in the absence of Ca^2+^ and the presence of increasing concentrations of CpxSC. The following concentrations of ^15^N-C_2_B and CpxSC (μM/μM) were used (from black to purple): 32/0, 30/10, 28/19, 26/28, 24/36, 20/53, 17/65, 12/88. The intensities of cross-peaks decreased as CpxSC was added because ^15^N-C_2_B was diluted and binding to CpxSC causes cross-peak broadening. Contour levels were adjusted to compensate for these decreased intensities, but some cross-peaks may not be visible even after these adjustments at the higher CpxSc concentrations. In addition some of the cross-peaks in the middle of the titration are not observed because of chemical exchange broadening. The red boxes indicate the cross-peaks that are displayed in the expansions of Fig. 4.

**Figure S13.**
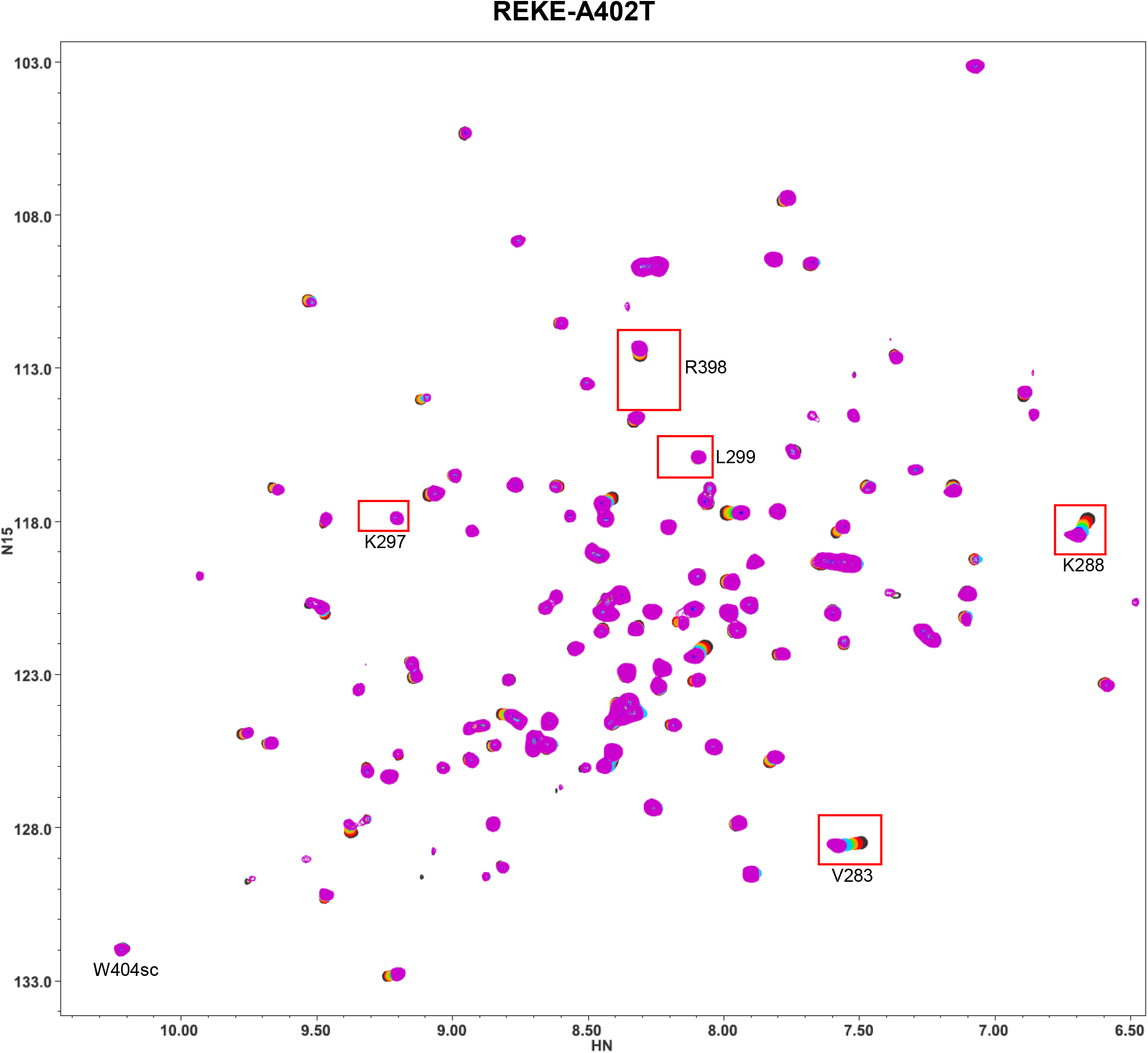
Superposition of ^1^H-^15^N TROSY-HSQC spectra of ^15^N-C_2_B domain REKE-A402T mutant in the absence of Ca^2+^ and the presence of increasing concentrations of CpxSC. The following concentrations of ^15^N-C_2_B and CpxSC (μM/μM) were used (from black to purple): 32/0, 30/10, 28/19, 26/28, 24/36, 20/53, 17/67, 12/88. The intensities of cross-peaks decreased as CpxSC was added because ^15^N-C_2_B was diluted and binding to CpxSC causes cross-peak broadening. Contour levels were adjusted to compensate for these decreased intensities, but some cross-peaks may not be visible even after these adjustments at the higher CpxSc concentrations. The red boxes indicate the cross-peaks that are displayed in the expansions of Fig. 4.

**Figure S14.**
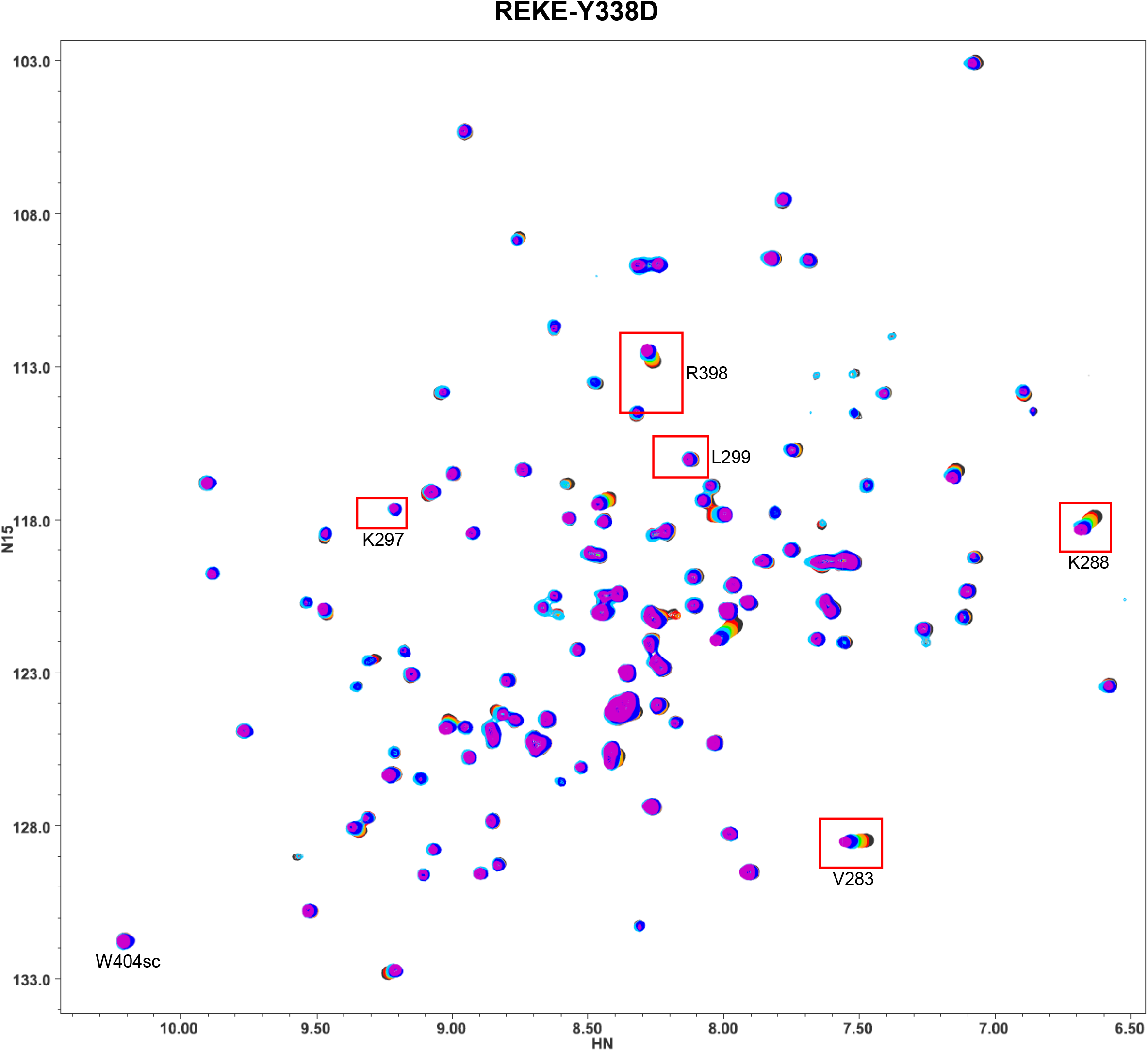
Superposition of ^1^H-^15^N TROSY-HSQC spectra of ^15^N-C_2_B domain REKE-Y338D mutant in the absence of Ca^2+^ and the presence of increasing concentrations of CpxSC. The following concentrations of ^15^N-C_2_B and CpxSC (μM/μM) were used (from black to purple): 32/0, 30/10, 28/19, 26/28, 24/36, 20/53, 17/68, 12/88. The intensities of cross-peaks decreased as CpxSC was added because ^15^N-C_2_B was diluted and binding to CpxSC causes cross-peak broadening. Contour levels were adjusted to compensate for these decreased intensities, but some cross-peaks may not be visible even after these adjustments at the higher CpxSc concentrations. The red boxes indicate the cross-peaks that are displayed in the expansions of Fig. 4.

**Figure S15.**
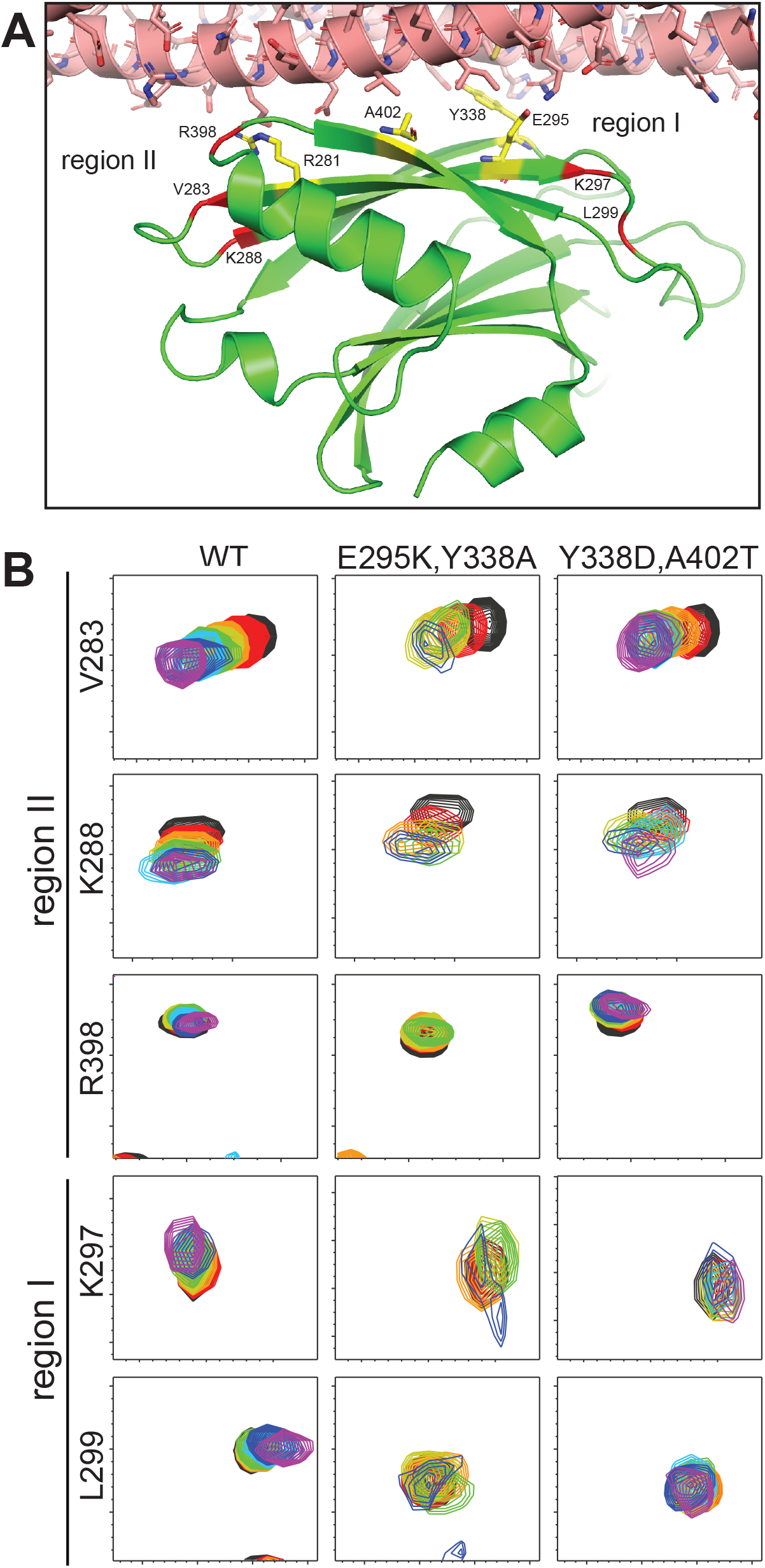
The Syt1 C_2_B domain still binds to the SNARE complex through region II of the primary interface when binding through region I is abolished. (*A*) Close-up view of the primary interface with the SNARE complex represented by a ribbon diagram and stick models (nitrogen atoms in dark blue, oxygen in red, sulfur in yellow orange and carbon in salmon color), and the Syt1 C_2_B domain represented by a green ribbon diagram and stick models for residues that were mutated in the NMR experiments of Fig. 4 and S6-S14 (carbon atoms in yellow). Residues corresponding to the cross-peaks shown in panel (*B*) and Fig. 4 are indicated in red on the ribbon diagram. These residues and those that were mutated are labeled. The diagram was generated with a crystal structure of a Syt1-SNARE complex (PDB accession number 5KJ7). (*B*) The diagrams show expansions of ^1^H-^15^N TROSY HSQC spectra of WT and mutant ^15^N-C_2_B domain (as indicated above) acquired in isolation (black contours) or increasing concentrations of CpxSC (rainbow colours). The residues corresponding to the cross-peaks shown in the expansions and the regions where they are located are indicated on the left. The following concentrations of ^15^N-C_2_B mutant and CpxSC (μM/μM) were used (from black to purple): WT 32/0, 30/10, 28/19, 26/28, 24/36, 20/51, 17/64, 12/85; E295K/Y338A 32/0, 30/11, 27/21, 25/30, 23/38, 19/59; Y338D/A402T 32/0, 30/10, 28/19, 26/28, 24/36, 20/51, 17/64, 12/86. The intensities of cross-peaks decreased as CpxSC was added because ^15^N-C_2_B was diluted and binding to CpxSC causes cross-peak broadening. Contour levels were adjusted to compensate for these decreased intensities.

**Figure S16.**
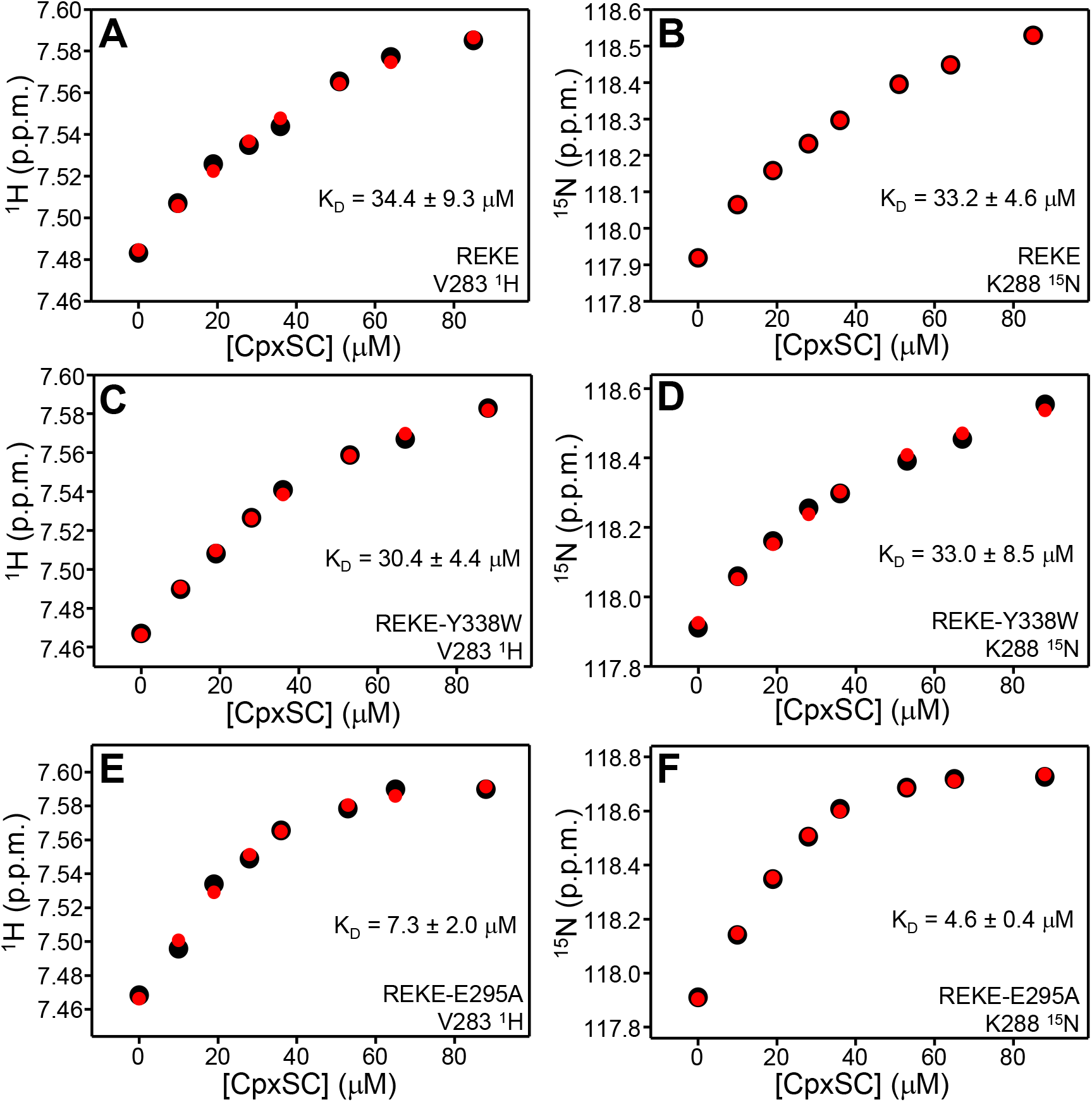
The E295A mutation enhances the affinity of the Syt1 C_2_B domain for the SNARE complex. (*A-F*) Plots of the ^1^H chemical shift of the V283 cross-peak (*A, C, E*) or the ^15^N chemical shift of the K288 cross-peak from REKE (*A, B*), REKE-Y338W (*C, D*) or REKE-E295A mutant ^15^N-C_2_B domain in the titrations with CpxSC monitored by ^1^H-^15^N TROSY-HSQC spectra (Fig. 4, S9, S11, S12). Note that the C_2_B domain concentration also changed during the titration as indicated in the corresponding figure legends. The black circles show the experimental data and the red circles show the predicted values after fitting the data to a single-site binding model using the equation f=d0+(df-d0)*(p+x+k-sqrt((p+x+k)^2-4*p*x))/(2*p), where f is the corresponding chemical shift, d0 is the chemical shift of free ^15^N-C_2_B domain, df is the chemical shift of CpxSC-saturated ^15^N-C_2_B domain, p is the ^15^N-C_2_B domain concentration, x is the CpxSC concentration and k is the K_D_ (all concentrations in μM units). The K_D_ values obtained for each graph are indicated together with the errors yielded by the fitting procedure.

**Figure S17.**
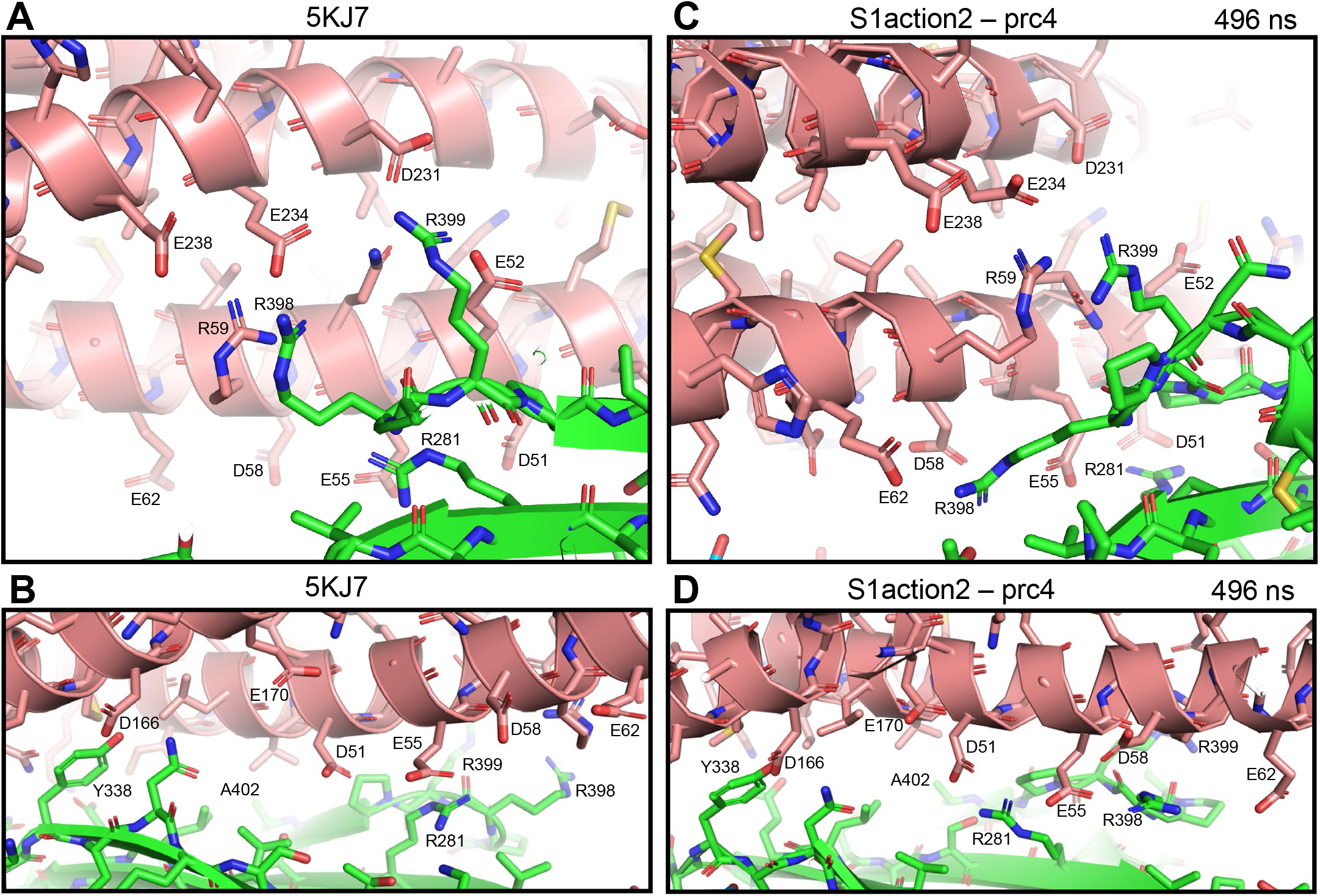
Diagrams illustrating differences observed in the primary interface of crystal structures of Syt1-SNARE complexes and in MD simulations. (*A, B*) Different views of region II a crystal structure of a Syt1-SNARE complex (PDB accession number 5KJ7). (*C, D*) Different views of region II of one of the primed complexes (prc4) at 496 ns of the s1action2 simulation. Proteins are represented by ribbon diagrams and stick models with nitrogen atoms in dark blue, oxygen in red, sulfur in yellow orange and carbon in salmon color (SNARE complex) or green (Syt1 C_2_B domain). Acidic residues from the SNAREs and the three arginines of the C_2_B domain are labeled. R59 of SNAP-25 is also labeled, as it may affect the interactions between C_2_B arginines and SNARE acidic residues.

**Figure S18.**
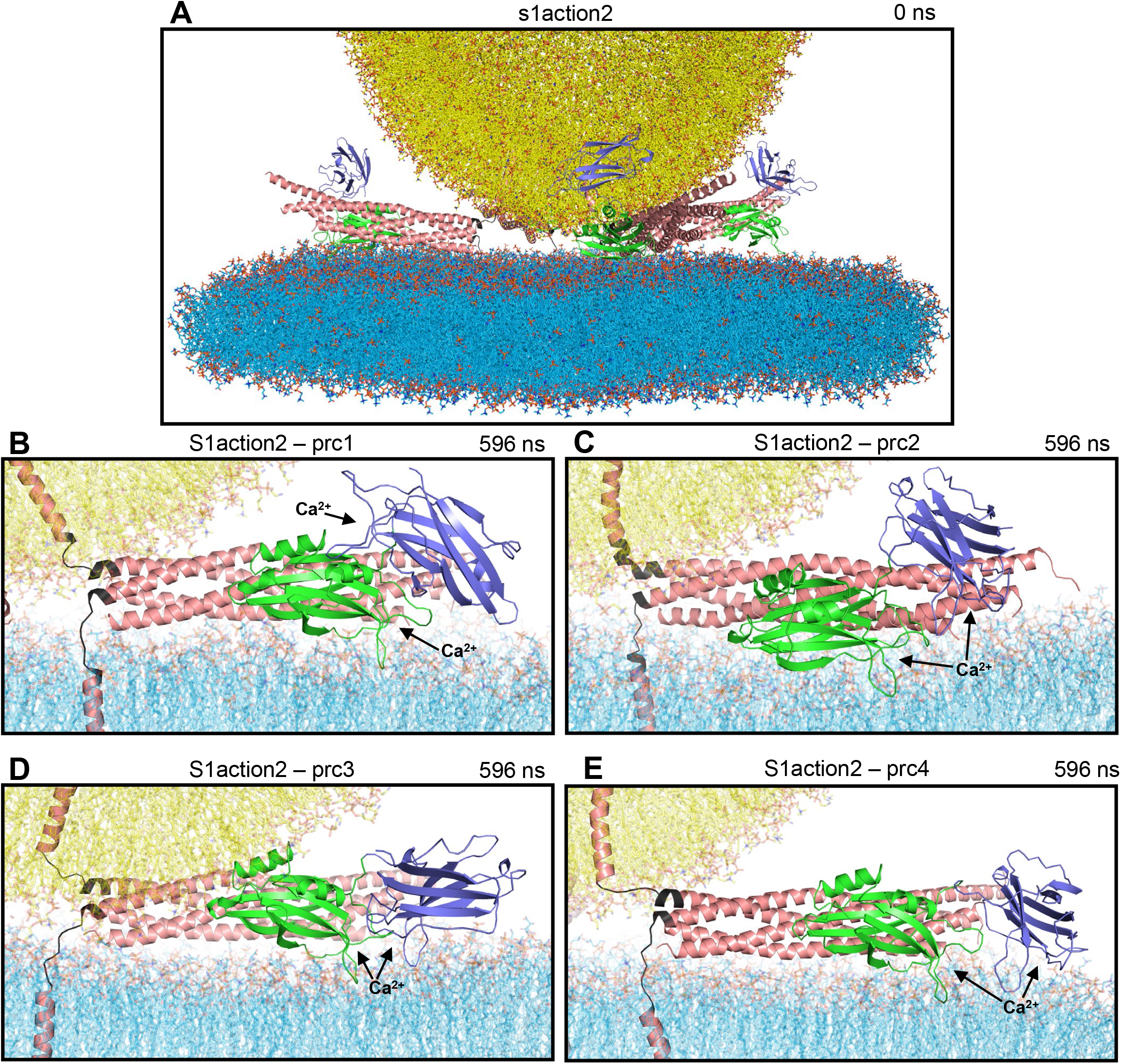
MD simulation of primed Syt1 C_2_AB-SNARE complexes bridging a vesicle and a flat bilayer (s1action2 simulation). (*A*) Initial configuration of the system. (*B-E*) Close-up views of the four primed complexes (prc1-prc4) after 596 ns of MD simulation. The arrows point at the Ca^2+^-binding sites of the C_2_A and C_2_B domains. Lipids are represented by stick models and proteins by ribbon diagrams with the same color coding as in Fig. 1, except that the jxt linkers of synaptobrevin and syntaxin-1 are colored in dark gray. Note the similarity of the orientations of the C_2_B domain with respect to the SNARE complex and the flat bilayer for the different complexes, the different conformations adopted by the jxt linkers, and the different locations of the C_2_A domain, although in three complexes (prc2-prc4) the Ca^2+^-binding loops of the C_2_A domain are located close to those of the C_2_B domain, which can facilitate a cooperative action of both domains.

**Figure S19.**
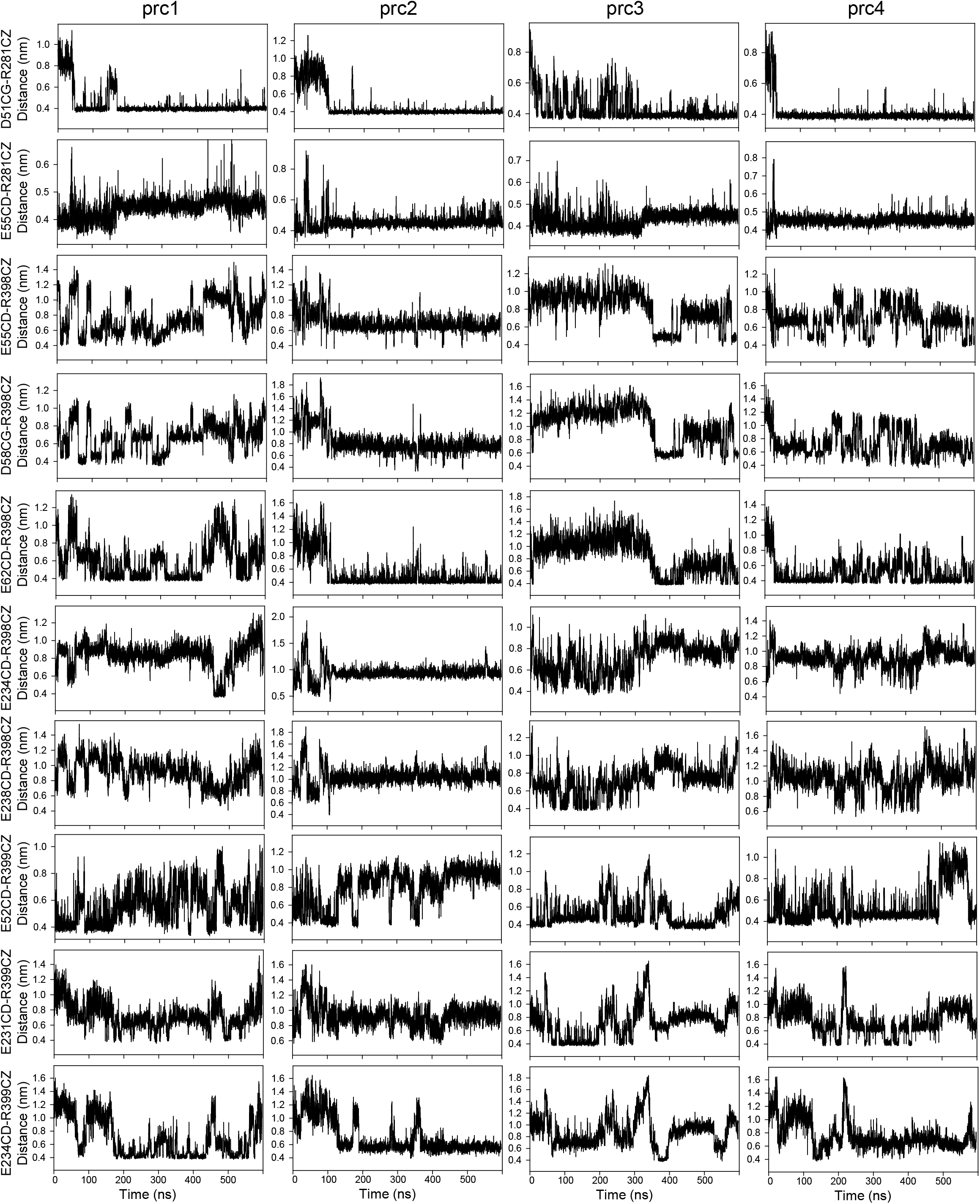
Distances between Syt1 C_2_B domain arginines and SNARE acidic residues at region II of the primary interface during the s1action2 MD simulation. The time dependence (0.1 ns steps) of the distances indicated at the y axes is shown for the four complexes (prc1-prc4) of the simulation.

**Figure S20.**
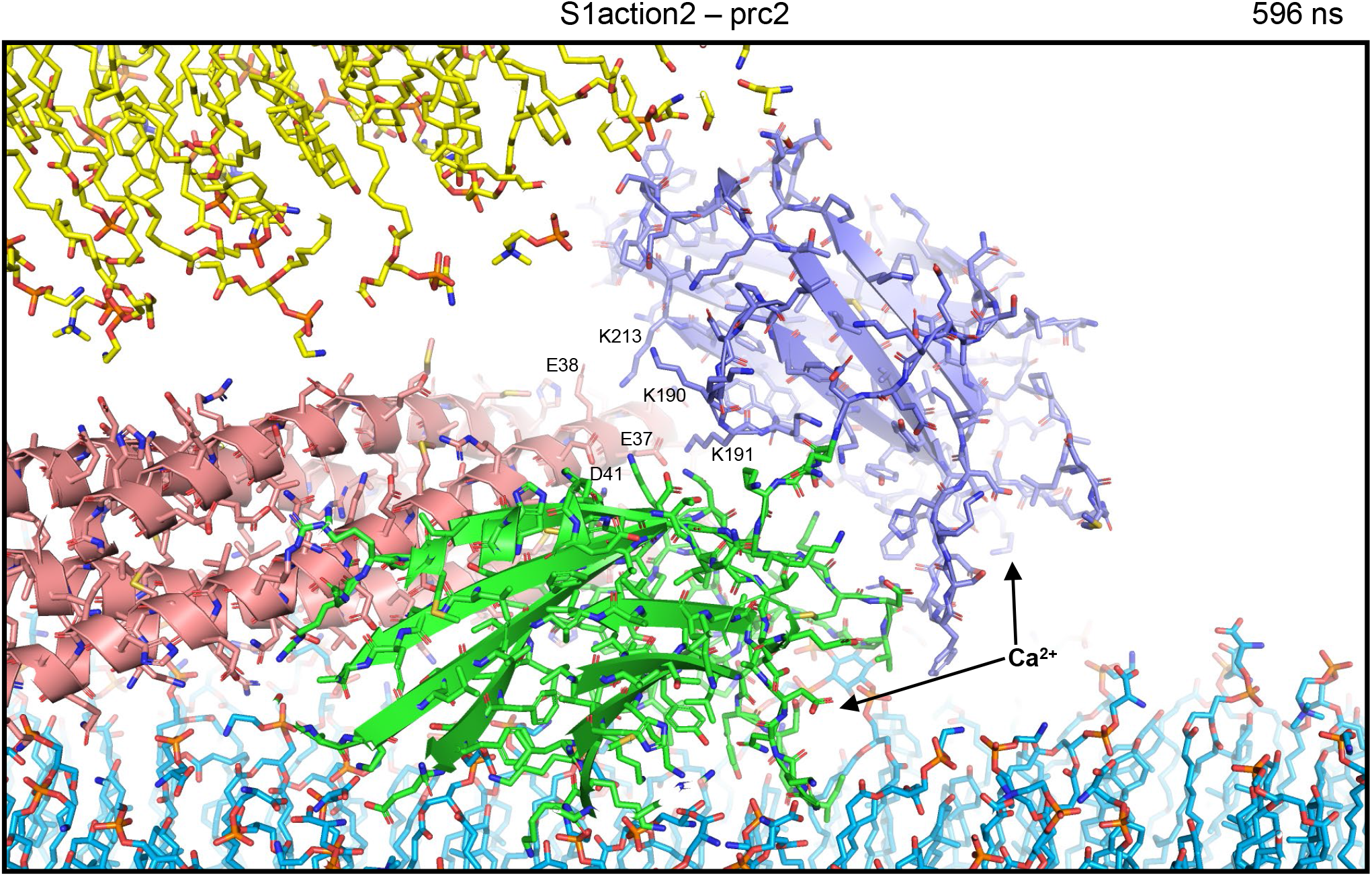
Close-up view of one of the primed complexes (prc4) of the s1action2 MD simulation at 596 ns. Lipids are represented by stick models and proteins by ribbon diagrams and stick models with the same color coding as in Fig. 1 and 2. Basic residues of the C_2_A domain that are close to an acidic patch of the SNARE complex are labeled. The arrows indicated the Ca^2+^-binding sites of the two C_2_ domains.

